# Mammalian octopus cells are direction selective to frequency sweeps by synaptic sequence detection

**DOI:** 10.1101/2021.11.29.470040

**Authors:** Hsin-Wei Lu, Philip Smith, Philip Joris

## Abstract

Octopus cells are remarkable projection neurons of the mammalian cochlear nucleus, with extremely fast membranes and wide frequency tuning. They are considered prime examples of coincidence detectors but are poorly characterized *in vivo*. We discover that octopus cells are selective to frequency sweep direction, a feature that is absent in their auditory nerve inputs. *In vivo* intracellular recordings reveal that direction selectivity does not derive from cross-channel coincidence detection but hinges on the amplitudes and activation sequence of auditory nerve inputs tuned to clusters of “hotspot” frequencies. A simple biophysical model of octopus cell excited with real nerve spike trains recreates direction selectivity through interaction of intrinsic membrane conductances with activation sequence of clustered inputs. We conclude that octopus cells are sequence detectors, sensitive to temporal patterns across cochlear frequency channels. The detection of sequences rather than coincidences is a much simpler but powerful operation to extract temporal information.

## Introduction

How single cells and small neural circuits extract sensory features via elemental computations is an important theme in systems neuroscience. Well-studied model systems include circuits producing tuning to visual orientation and direction (Ferster and Miller, 2000; Hubel and Wiesel, 1974; Vaney et al., 2012), to whisker deflection (Simons, 1978), and to binaural disparities (Yin et al., 2019). In hearing, many elemental computations have a temporal basis, often hypothesized to be instantiated by coincidence detection combined with time delays. Examples are the processing of interaural time delays (Joris and van der Heijden, 2019) and enhanced temporal coding in the cochlear nucleus (Joris and Smith, 2008; Joris et al., 1994). Coincidence detection is a simple concept dovetailing quite naturally with basic neural properties and is easily implemented computationally, but physiological evidence is fragmentary.

Octopus cells of the mammalian cochlear nucleus are named for the unique morphology of their dendrites, oriented in a common direction across the tonotopic array of auditory nerve fibers (ANFs) (Harrison and Irving, 1966; Osen, 1969). Octopus cells have very leaky membranes (input resistance <10MΩ) owing to abundant low-voltage activated potassium channels (KL) and hyperpolarization-activated cation channels (HCN) (Bal and Oertel, 2000, 2001). This results in tiny EPSPs and action potentials, ultrafast membrane time constant and narrow integration time window (∼1ms) (Cao and Oertel, 2011; Golding et al., 1995, 1999; McGinley and Oertel, 2006), leading to the hypothesis that coincident activation of ANFs from a wide cochlear sector is required to achieve sufficiently fast and large depolarization to initiate spiking. Their status as monaural coincidence detectors inspired many computational studies (Cai et al., 1997, 2000; Hemmert et al., 2005; Kalluri and Delgutte, 2003a, 2003b; Kipke and Levy, 1997; Levy and Kipke, 1997; Manis and Campagnola, 2018; McGinley et al., 2012; Rebhan and Leibold, 2021; Spencer et al., 2012) but there is a dearth of *in vivo* data because the membrane properties hamper such recordings.

A remarkable finding is that octopus cells entrain to transient sounds like clicks by firing exactly one spike per click with submillisecond precision, even for trains >500Hz (Godfrey et al., 1975; Oertel et al., 2000; Pfeiffer, 1966). Clicks trigger a dispersive cochlear traveling wave from high (basal) to low (apical) characteristic frequency (CF) regions causing an asynchronous activation of ANFs across CFs, spanning milliseconds. How can an octopus cell, with ∼1ms integration time window, respond to such a temporally smeared input? A computational model supports the hypothesis that the peculiar dendritic morphology of octopus cells compensates for cochlear delay (McGinley et al., 2012). Simply put, delayed (low-CF) inputs innervate soma and proximal dendrites, while high-CF inputs innervate distal dendrites. As a result, EPSPs triggered by these various inputs coincide near the spike generator and nullify cochlear dispersion.

We used frequency-modulated (FM) stimuli to test the hypothesis of cochlear delay compensation and discovered a new form of tuning which unexpectedly shows wide tolerance for delays across frequency. Further investigation into the underlying mechanisms revealed that the critical process is not coincidence detection across frequency channels, but a sensitivity to the sequence of activation of discrete clustered inputs to these neurons. The simplicity of the mechanism suggests it has wider relevance in the creation of temporally based tuning properties which are traditionally thought to reflect coincidence detection.

## Results

### Gerbil Oi cells fit classical description of octopus cells

Because octopus cells have been rarely studied in gerbil (Feng et al., 1994; Ostapoff et al., 1994), we examined general response properties using axonal recordings with sharp pipettes (Smith et al., 2005) to bypass the tiny somatic action potentials (Golding et al., 1995). Two types of onset response, Oi and OL (see Methods), have been associated with octopus cells, where Oi but not OL neurons lack sustained firing to pure tones. Our sharp recordings yielded 33 Oi and 7 OL neurons (Figure 1A,B). A criterion of sustained rate to tones of 30 spikes/s (Figure 1B) resulted in a segregation of the two groups along several dimensions. Frequency tuning curves typically showed high thresholds in Oi neurons but were more nerve-like in shape and threshold in OL neurons (Figure 1C). Examples of both types were seen in a single animal indicating high threshold was not due to poor cochlear condition. Click trains evoked sustained firing in Oi neurons with exquisite temporal locking and entrainment (1 spike/click) up to 500-600 Hz and quickly declining firing rates for further increases in click frequency (Figure 1D,E). Temporal locking to clicks was extremely precise near 100 Hz (vector strengths >0.999) and remained high (>0.5) sometimes up to 1kHz (Figure 1F,G). Entrainment was more limited in OL neurons, with higher firing rates (Figure 1D,E) but lower vector strengths (Figure 1F,G) at high click frequencies. The results from the sharp recordings were corroborated by intra- and extracellular recordings (Figure 1B-F, solid lines, symbol legend in B). Properties of Oi but not OL units thus fit those of labeled octopus cells in cat (Oertel et al., 2000; Smith et al., 2005). Our labeling results (not shown) also confirmed that Oi cells exhibit the morphology and anatomical location of octopus cells as previously described. We therefore focus on Oi neurons in subsequent sections.

**Figure 1.**
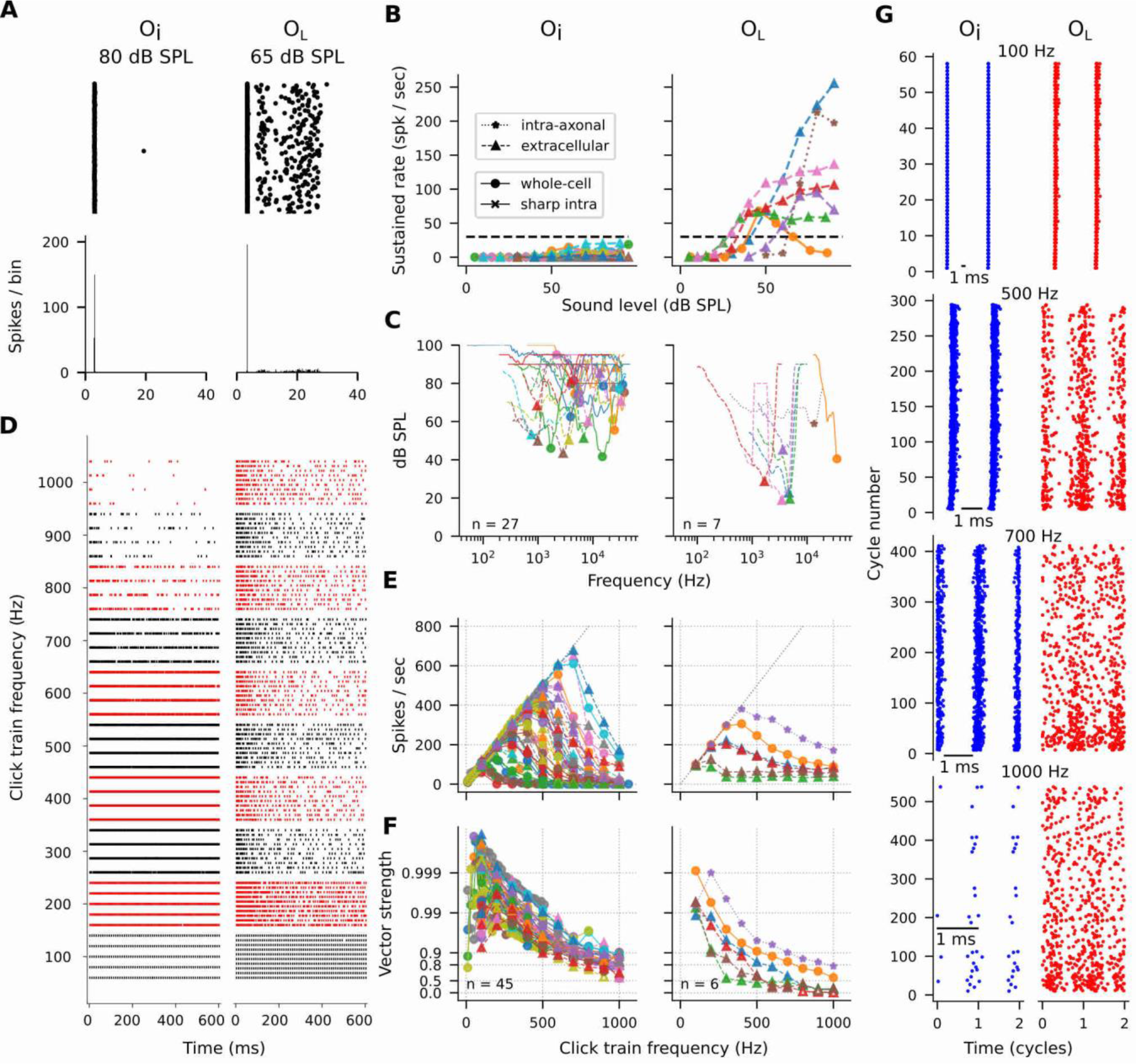
Gerbil Oi neurons fit classical description of octopus cells. Panels (A,D,G) show individual examples; in panels (B,C,E,F) each line represents one cell. **(A)** Oi cell (left, CF: 23.9 kHz) and OL cell (right, CF: 4.8 kHz) distinguished by their response to short (25 ms) tones. Top and bottom show dotrasters and PSTHs, respectively. **(B)** Sustained rate (measured between 10 – 25 ms) as a function of tone level. The horizontal dashed line shows the Oi vs. OL criterion (30 spikes/s). **(C)** Threshold tuning curves. Minimal thresholds were significantly higher in Oi than OL cells (Oi: 67.6 ± 14.4 dB SPL, n = 27; OL: 33.4 ± 14.1 dB SPL, n = 7; p < 0.0001, t-test). Colors consistent with (B). **(D)** Dotrasters of responses to click trains of increasing frequency; same cells as in A. **(E and F)** Population plots of spike rate (E) and vector strength (F) vs. click train frequency for Oi (left) and OL (right) cells. The unity line indicates perfect entrainment (firing rate = click train frequency). Open symbols in (F) indicate statistically non-significant points (Rayleigh test > 0.001). All Oi units, but only 2 / 6 of the OL units, had vector strength > 0.9 for 500-Hz click trains. Colors consistent across (E) and (F). **(G)** Cycle-based dotrasters to click trains from 100 - 1000 Hz for the Oi (blue) and OL (red) cell in (D). In each raster, the response to the earliest cycles is shown at the bottom and to the latest cycles at the top, and the raster is plotted twice (two cycles) for easier visualization. The Oi cell shows temporally precise responses but often skips cycles at the highest frequencies, while the OL cell gives a sustained response at all frequencies with more limited temporal resolution.

### Octopus cells, but not nerve fibers, entrain to frequency sweeps with directional tuning

The highly entrained click train responses are not a strong test of the cochlear delay compensation hypothesis because delay across frequencies is not varied. We parametrically varied this delay using Schroeder harmonic complexes (Schroeder, 1970). These stimuli consist of a series of equal-amplitude harmonic tones (Figure S1B), whose phase relation is systematically manipulated (parameter C, see Methods) to generate repetitive frequency sweeps of different direction and speed (Figure S1A). The waveform is a click train for C=0, and for C≠0 is an upward (C<0) or downward (C>0) FM sweep. The sweep duration shortens as C approaches zero. If dendritic compensation of cochlear delay is critical, only click trains (C=0) should provide sufficient coincidences for entraining responses and any deviation from C=0 should result in a reduced response.

We found that octopus cells entrain not only to click trains (C=0), but also to non-zero C values that produce large delays across frequencies. Figure 2 shows two representative examples. One neuron (Figure 2A,D,E) responds strongly to C≥0. It entrains not only to C=0 but also to C=0.5, which generates a frequency sweep (30 to 0.2 kHz) spread over a time span (2.5 ms) comparable to the cochlear delay (Figure 2A). The other neuron (Figure 2C,F,G) responds strongly to C≤0. It entrains to both click trains (C=0) and a long frequency sweep (10 ms, from 0.05 to 10 kHz at C=-0.5). These results suggest that large delays, imposed by FM sweeps across frequency channels, do not necessarily cause a precipitous drop in firing. This result questions the importance of coincidences across frequency and of precise dendritic compensation for cochlear delay.

**Figure 2.**
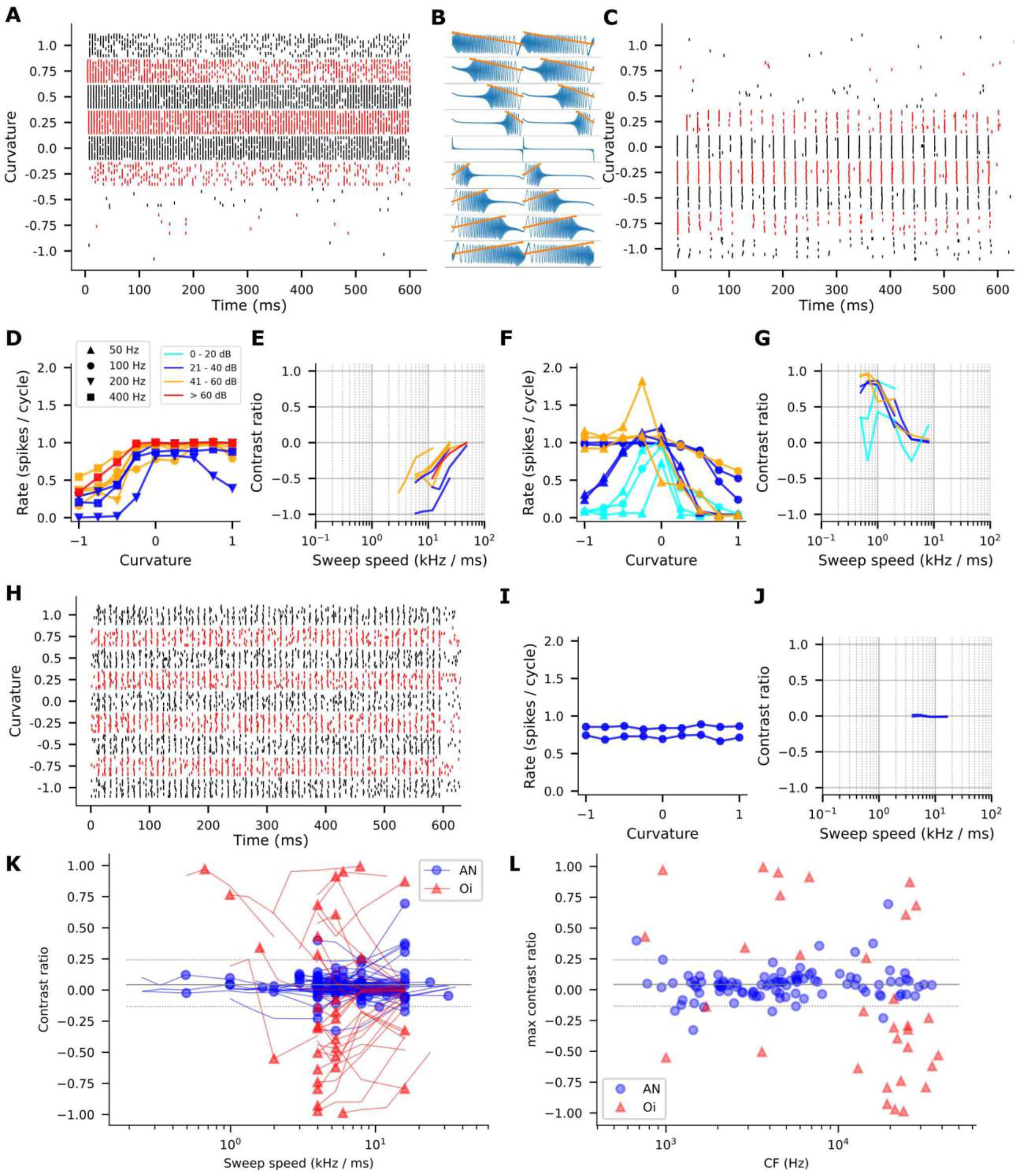
Octopus cells but not ANFs show rate tuning to the sweep direction of Schroeder stimuli. **(A and C)** Spike dotrasters of an octopus cell (A, CF: 24 kHz) tuned to downward sweeps (C > 0) and the other (C, CF: 0.96 kHz) tuned to upward sweeps (C < 0). Schroeder parameters for (A): F0=200Hz, 25 dB SPL, harmonics #1 - #150; parameters for (C): F0=50Hz, 40 dB SPL, harmonics #1 - #100. **(B)** Generic stimulus waveforms at C values corresponding to responses in (A) and (C). Orange: instantaneous frequency. **(D-G)** Rate curves and contrast ratios at various conditions from the cell in (A) and (C). (D, E) are the curves from cell in (A); (F, G) from cell in (C). Note for each cell, the rate tuning is largely consistent across F0s and sound intensity, i.e. (D, E) favors downward sweeps (C > 0) and and (F, G) favors upward sweeps (C < 0). At the lowest intensities (F, cyan traces), firing rate peaked at C=0, however as intensity increases, firing rate grows asymmetrically with a preference for upward sweeps (C < 0). **(H-J)** Dotraster, rate curve, and contrast ratios for an ANF with CF (0.95 kHz) similar to the octopus cell in (C). There is no rate tuning (I) and contrast ratios are near 0 (J) for two sound intensities tested. **(K)** Population plots of the contrast ratio vs. sweep speed in ANFs (AN, n=89) vs octopus cells (Oi, n=32). Each line represents the dataset (i.e. responses to a full set of Schroeder stimuli at one SPL and one F0) that contains the largest contrast ratio (solid circles) for that cell. Horizontal lines indicate the median value (0.04, solid) and 5 - 95% range (-0.13 – 0.24, dotted) of the largest contrast ratios among ANFs. Most octopus cells have their largest contrast ratios outside this range. **(L)** Population plots of the maximal contrast ratio vs CF. Symbols and horizontal lines are as in (K). There is a trend for high-CF octopus cells to prefer downward sweeps.

The different tuning of the two neurons (to downward sweeps, C>0, in Figure 2A, and to upward sweeps, C<0, in Figure 2C) persists for a range of SPL and F0 (Figure 2 D,F), although it tends to decrease with intensity. For example, the cell in Figure 2F responded maximally to C=0 at the lowest intensities. Nevertheless, the results reveal a form of tuning which can be equivalently described as direction selectivity to frequency sweeps or monaural phase sensitivity.

We quantified the sign and magnitude of direction selectivity with a conventional contrast ratio for each sweep speed (i.e. each pair of C values, e.g. ±0.5). Positive ratios indicate higher firing rate for upward sweeps, negative for downward sweeps, and zero for neither direction (see Methods). We found that the sign of direction selectivity is quite invariant across intensity and F0. For the two representative cells, nearly all contrast ratios were ≥ 0 for one cell and ≤ 0 for the other (Figure 2 E,G). The curves tend to converge to 0 as sweep speed increases: this reflects the finding that responses tend to be similar across small C values (Figure 2D,F) and show the largest differences usually for pairs with slow sweep speeds (C= ±1), which impose large delays across frequency channels.

Octopus cells do not simply inherit their direction selectivity from their inputs. ANFs show no changes in firing rate for up vs. downward sweeps (Figure 2H-J). The difference in the direction selectivity between ANFs and octopus cells is evident at the population level. Figure 2K shows contrast ratios from 89 ANFs and 32 octopus cells as a function of sweep speed. Almost all octopus cells (97 %; 31/32) had their largest contrast ratios outside the 5-95% range of contrast ratios of ANFs (Figure 2K: horizontal dotted lines). Figure 2L shows maximal contrast ratios as a function of CF. At all CFs, octopus cells show marked direction selectivity.

### Frequency sweeps cause large differences in temporal pattern across the auditory nerve

The selectivity of octopus cells must arise from aspects of the ANF response other than their spike rates. Previous work reported a somewhat peakier temporal response to downward vs upward Schroeder sweeps, both at the cochlear mechanical level and for individual ANFs and certain cochlear nucleus neurons (Recio, 2001; Recio and Rhode, 2000; Summers et al., 2003). Figure 3A shows dotraster and cycle histograms from one ANF. The distribution of spikes is narrow for C=0 and broadens with increasing |C|. This is not due to drift in spike timing during repetitive stimulation: the timing is consistent across successive cycles (bottom vs. top of dotrasters). There is indeed an asymmetry with slightly narrower and taller cycle histograms for C>0 than for C<0. To examine whether this asymmetry can explain the direction selectivity of octopus cells (Figure 2K,L), we used an analysis based on the popular notion of octopus cells being coincidence detectors (Golding et al., 1995; Kalluri and Delgutte, 2003a).

**Figure 3.**
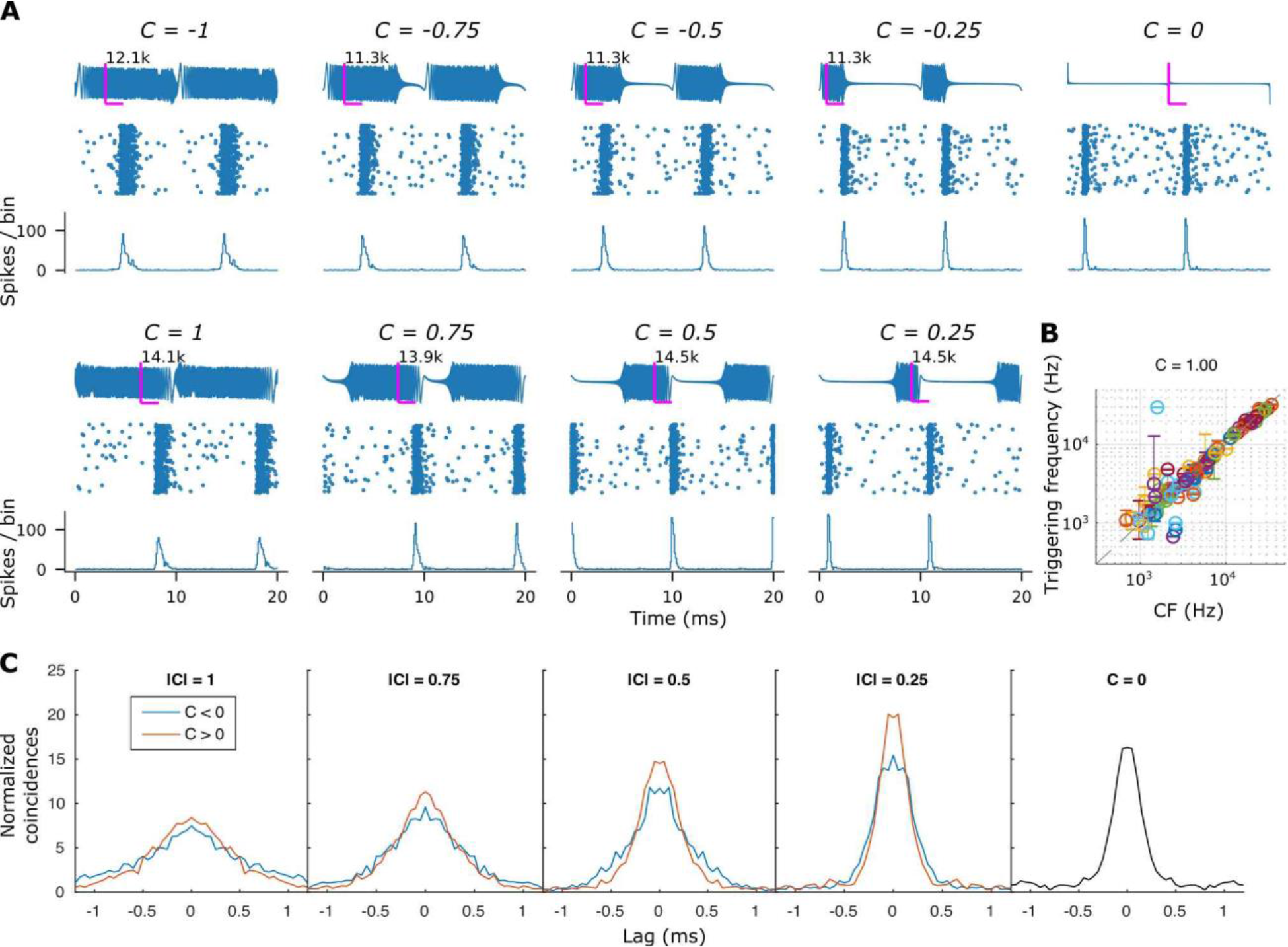
Temporal responses of single ANFs to Schroeder stimuli. **(A)** Cycle-based responses from one ANF (CF = 12.8 kHz, SR = 24 spikes / sec) to Schroeder stimuli (F0: 100 Hz, harmonics: #1 - #400, 30 dB SPL) at different curvatures. For each C, two cycles of the Schroeder waveform are shown above the dotraster and cycle histogram. Spike-triggering frequency can be estimated by using the spike latency to the click stimuli (C = 0), as indicated by the magenta lines and the numbers in each condition. Note that the triggering frequency is similar across different C values and is close to the CF (12.8 kHz). **(B)** Population plot of the spike triggering frequency in Schroeder stimuli (C = 1) vs CF of the ANFs. Each dot represents one cell (n = 103) and indicates the median value of the estimated triggering frequency. Error bars represents 1st - 3rd quartile. Note the dots lie mostly on the identity line (dashed), indicating the triggering frequency is close to the CF of the cell. **(C)** Shuffled autocorrelograms of the responses in (A), superimposed for C < 0 (blue) and C > 0 (red), show that the number expected coincidences is similar for stimuli with opposite sweep direction. In this cell, peak maxima are slightly higher for downward sweeps (C > 0, red).

We counted the number of coincidences across repetitions of different C values. Figure 3C shows the central peak of shuffled autocorrelograms (Joris, 2003) calculated for the responses shown in Figure 3A (Figure S2C shows additional examples), superimposed for opposing directions at the same speed. There are more coincident spikes (higher peaks) for the downward (C>0) stimuli. In some ANFs, the opposite pattern is observed (Figure S2C, rows 3 and 4). From the peak number of coincident spikes for opposite C values, we calculated contrast ratios. Although the resulting pattern shows a trend of higher contrast ratios at low CF (∼1kHz), the contrast ratios for higher CFs are small and cannot account for the pattern seen in octopus cells (Figure S2A,B).

For a given ANF, there are obvious latency differences between responses to various C-values. For example, for the ANF in Figure 3A, the response to the downward sweep is delayed relative to the upward sweep (C=1 vs -1). This is expected given its CF (12.8 kHz): a linear upward sweep from 100 Hz to 40 kHz traverses the tuning curve earlier than a downward sweep. Assuming a fixed delay between triggering sound and spiking, we can estimate the instantaneous frequency that triggers spikes (Heil et al., 1992; Nelken and Versnel, 2000). We used response latency (peak of the cycle histogram) to clicks (C=0) to estimate the triggering frequency in other conditions (magenta lines, Figure 3A). For the ANF population studied, this frequency is close to CF and is largely similar across sweeps at different speeds or directions (Figure 3B, Figure S3). Only for CFs below a few kHz, where the linear sweep passes the CF range very quickly and responses are generally more smeared even to transient stimuli, does the relationship become noisier. Given the bias of octopus cells to high CFs (Figure 1C, 2L, 6A2), this analysis provides an important means to derive CF from inputs to octopus cells (next Section).

Because the timing of instantaneous frequency changes with speed and direction of the Schroeder stimuli (Figure S1), the temporal pattern across ANFs tuned to different CFs changes accordingly. Figure 4A overlays cycle histograms from responses of 67 ANFs ordered with CF. For C=0, all frequencies start simultaneously (black vertical line) and cochlear delay is the only source of latency differences. The increase in latency with decreasing CF of responses to C=0 matches those to a 100-Hz click train (Figure 4B). For both stimuli, ANFs tuned to low CFs (<4kHz) have delayed responses compared to high-CF fibers. For C≠0, there is a frequency- dependent delay in the stimulus itself which generates a curved pattern (Figure 4A: black lines). As expected, the population response pattern follows the timing of the instantaneous frequencies in the stimulus combined with cochlear delay: this enlarges the range of delays for downward FM sweeps (C>0, lower row of Figure 4A) and reduces the range for upward sweeps (C<0). However, combining the observed bias of octopus cells to high frequencies (Figures 1C,2L,6A2 and Godfrey et al., 1975) with the absence of a measurable cochlear delay for CF>4 kHz (Figure 4, see also Huet et al. 2016), the overall effect of cochlear delay is unlikely to be a significant factor in the sensitivity to FM sweeps. These results strongly suggest that the source of directional tuning of octopus cells is in the delay patterns across ANFs evoked by Schroeder stimuli.

**Figure 4.**
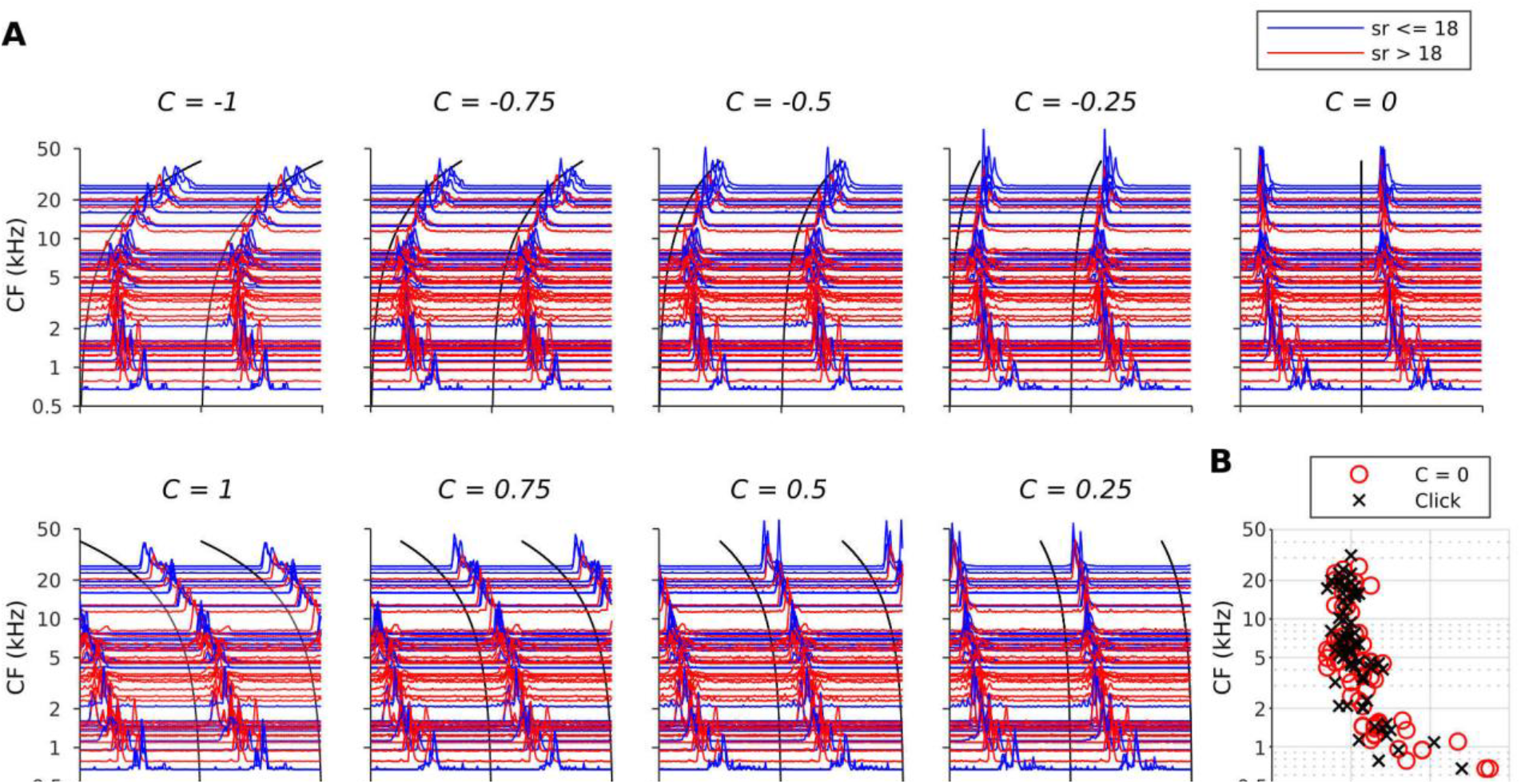
Schroeder stimuli evoke different delay patterns across frequencies in the ANF. **(A)** Population waterfall plots of ANF cycle histograms, sorted by CF (note the logarithmic ordinate), in response to Schroeder stimuli (F0: 100 Hz, harmonics: #1 - #400, 30 dB SPL). Two cycles are presented for clarity. The trajectory of instantaneous frequency of the stimuli is superimposed on the cycle histograms (black curves). ANFs with low- and high-spontaneous rate (sr <= 18 vs > 18) are color coded in blue and red, respectively (n = 60). **(B)** Peak latency of the responses to C = 0 (red circles). For comparison, the latency of responses to click trains (100 Hz, 70 dB SPL) are also shown (n = 67). A clear cochlear delay is seen in CFs < 4 kHz in both conditions.

### Spikes in octopus cells can be triggered by different instantaneous frequencies depending on sweep rate and direction

To understand how FM sweeps trigger entrained spikes in octopus cells, we first examined the temporal relation between spiking and instantaneous frequency, using the latency-based technique previously validated on ANFs (Figure 3A,B). Given the brief EPSP integration time window (∼1ms) (Ferragamo and Oertel, 2002; McGinley and Oertel, 2006), spike timing should reveal the spike-triggering frequency across different stimulus conditions. Figure 5A,B shows responses from a cell with low response rates at the most negative C values. Surprisingly, the triggering frequency differs for downward sweeps (C>0, ∼20 kHz) versus upward sweeps (C<0, ∼14 kHz), while both frequencies re-emerge at the most negative C values as two clusters of spiking. Figure S4 shows a similar pattern for a different cell, with responses triggered near 17 kHz for upward and 23 kHz for downward sweeps and 3 modes per cycle at the lowest response rates. The responses of Figure 5C reveal an even more complex pattern. The tuning to Schroeder stimuli in this cell is unusual in that it gives the lowest response to C values near 0. Superb phase-locking is present at all C values. For values |C|≥0.5, there are multiple triggering frequencies and high firing rates: a response is present for triggering frequencies near CF (∼20 kHz), followed by a response with a triggering frequency of only a few kHz. For upward sweeps, there is an opposite pattern.

**Figure 5.**
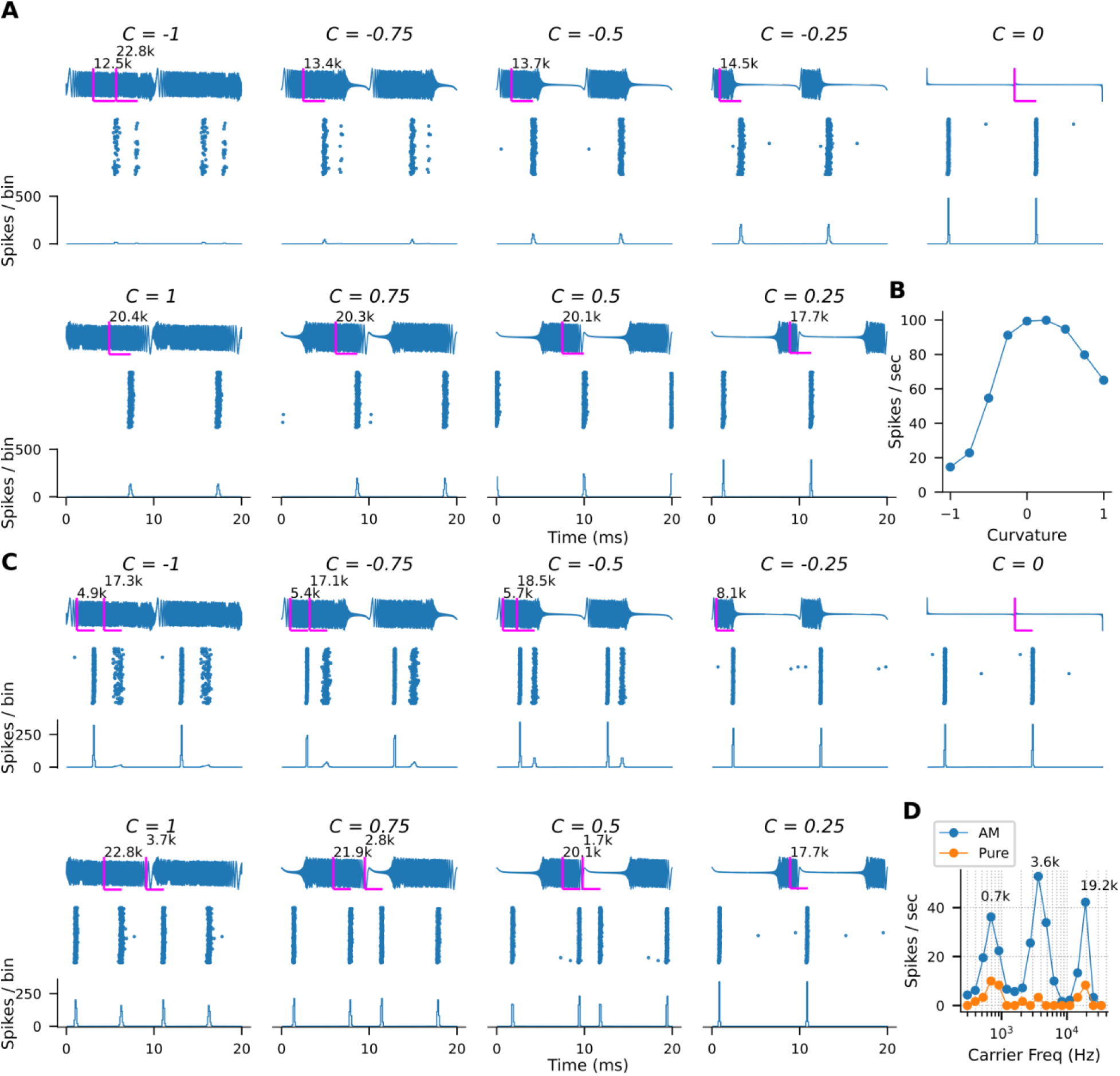
Octopus cells phase lock to instantaneous frequency in Schroeder stimuli. **(A)** Cycle-based dotrasters and cycle histograms of responses to Schroeder stimuli (F0: 100 Hz, harmonics: #1 - #400, 30 dB SPL) from a representative octopus cell favoring downward sweeps (C > 0). The spikes strongly phase lock to the frequency sweep. As in Figure 4, the triggering frequency of the spike can be estimated using the latency to C = 0, and is indicated in each panel. For multimodal responses more than one triggering frequency is obtained. **(B)** Rate vs Curvature relation of the response in (A). The cell fired at higher rate to downward sweeps (C > 0). **(C)** Example response from another octopus cell. Same stimulus parameters as in (A) but at 50 dB SPL. The cell often shows responses to two triggering frequency ranges (one at 18-22 kHz and the other at 2 – 5 kHz), which match the two peaks in the response area in (D). **(D)** Iso-intensity contours of the cell in (C) to pure (orange) and AM (blue) tones at 70 dB SPL. Peak responses at carrier frequencies of 3.6 and 19.2 kHz match the triggering frequencies in (C).

Multimodal responses as in Figures 5 and S4 are not consistent with the general assumption that octopus cells receive input from a broad and continuous sector of the cochlea and fire when these inputs are coincident. Rather, these results suggest the existence of “hotspots” of different CF and threshold in the array of ANFs converging on a cell. The latency-based derivation of instantaneous frequencies triggering a response is, however, an indirect assessment. When recording time allowed, we obtained a more direct, suprathreshold estimate by stepping pure and/or amplitude-modulated (AM, see Methods) tones of fixed intensity across a wide frequency range. Figure 5D shows responses to pure and AM tones at 70 dB SPL. Consistent with previous reports (Recio-Spinoso and Rhode, 2020; Rhode, 1994), AM stimuli evoke much higher sustained spike responses than pure tones. Remarkably, the responses area to AM tones reveals three discontinuous regions, and the carrier frequencies triggering the responses largely match the triggering frequencies in responses to Schroeder stimuli. Hence, these two independent measures suggest that inputs to octopus cells converge from a wide cochlear sector but are not smoothly distributed across frequencies.

### Intracellular recordings reveal “hotspots” in the inputs to a majority of octopus cells

The observations on the spike output to Schroeder stimuli are hard to reconcile with a hypothesis of coincidence detection across inputs with different CFs. We obtained *in vivo* intracellular responses from 16 octopus cells (12 with sharp electrodes, 4 whole-cell recordings) to Schroeder to examine how subthreshold responses are transformed into spikes. Figure 6A,B shows the median membrane potential of 15 representative cells in response to Schroeder stimuli, stacked on top of each other for 5 C values. A clear trend is observed: the EPSP was narrowest with the highest amplitude at C=0 and became broader as |C| moves towards 1. This suggests that inputs align maximally for the click-like stimulus, consistent with the input patterns measured in the ANFs (Figure 5).

**Figure 6.**
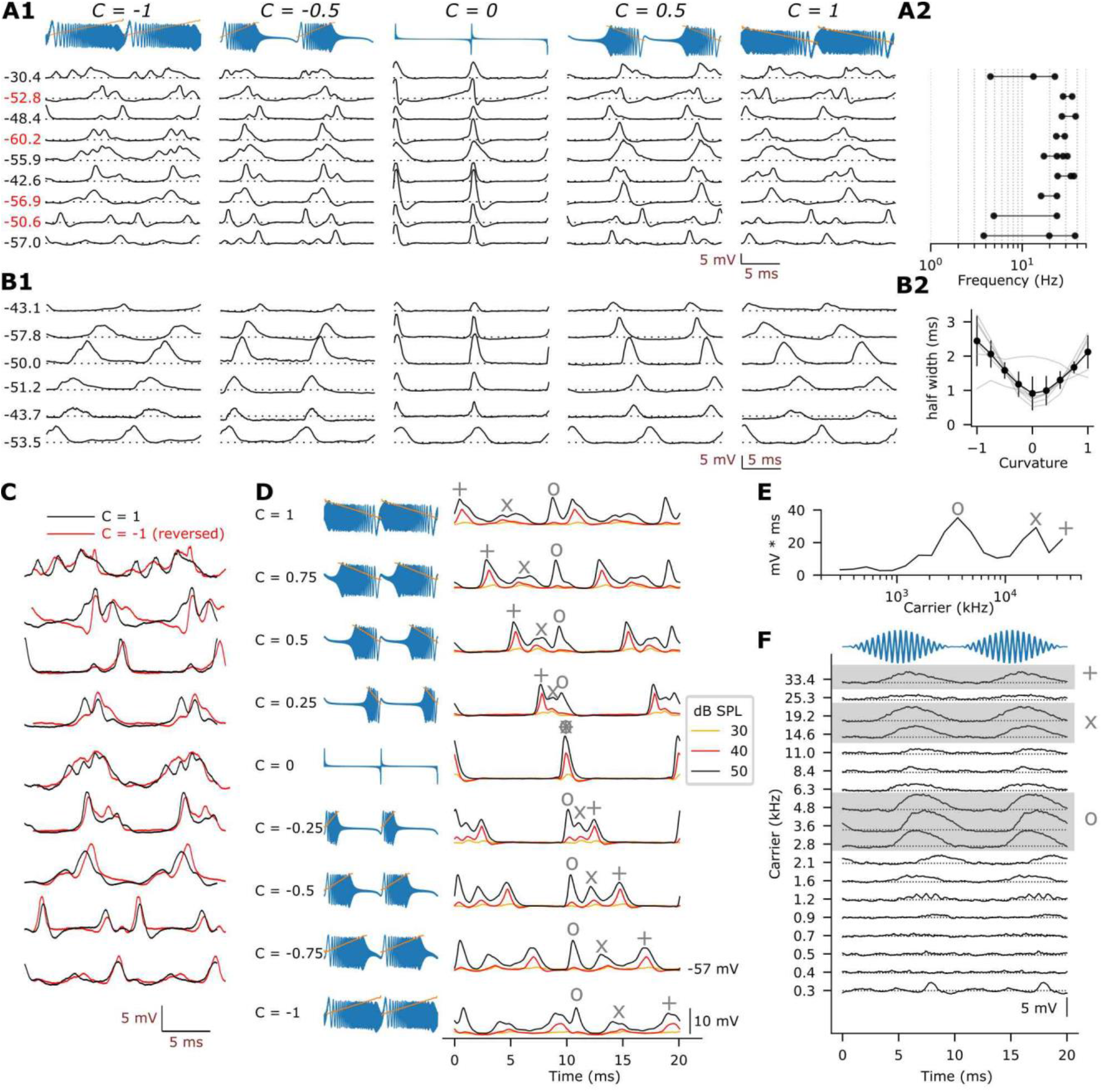
Intracellular recordings reveal “humps” in the response of octopus cells to frequency sweeps. **(A1)** Cycle-based median membrane potentials of 9 octopus cells in response to Schroeder stimuli (F0=100Hz, harmonic #1 - #400). Each column shows two response cycles for different phase curvatures. Dotted lines indicate the resting membrane potential, whose value is indicated on the left of the trace. Red fonts indicate recordings via whole-cell, others via sharp. There are clear “humps” in response to non-zero Schroeder stimuli, which we interpret as discrete EPSPs corresponding to hotspots in frequency tuning. As C approaches zero, the overall duration of the response narrows. **(A2)** Triggering frequencies of the “hump EPSPs” for responses in A1. **(B1)** Similar to (A1), but for 6 cells which did not show clear EPSP “humps”. **(B2)** The halfwidth of the EPSP narrows as C approaches 0. Gray, data from individual cells in B1; black, mean ± SD of the group in B1. **(C)** Time-reversed responses to C = -1 overlap well with responses to C = 1, suggesting that these humps are triggered by the same instantaneous frequencies. Traces are same as in A1. **(D-F)** An example cell where the hump-triggering frequencies in Schroeder stimuli (D) could be compared to hotspots in the response to AM tones (E, F). (D) shows the cycle-based median EPSP responses at different C values. The symbols (o, x, +) indicate the time points where the stimulus instantaneous frequency equals the triggering frequencies at 4, 19, 33 kHz: these time points correspond well with the EPSP peaks. Note that at as sound level increases from 40 to 50 dB SPL, the amplitude of EPSPs triggered by 4 and 19 kHz (o, x) increases much more significantly than the one triggered by 33 kHz (+), suggesting that the corresponding inputs to this cell have different thresholds. (E) shows the membrane potential integrals across carrier frequencies in the responses to AM stimuli (modulation frequency / depth: 100 Hz / 100%, 70 dB SPL) with same symbols as in D indicating the peak frequencies. (F) shows individual membrane potential traces (cycle-averaged, 2 cycles shown) to each carrier frequency: the triggering frequencies in (E) and (F) match those in (D), as indicated by the symbols. Dotted line in each trace indicates resting Vm at -57.8 mV. Blue trace on top: exemplar AM waveform.

Interestingly, in most cells (62.5%, 10/16), the broader EPSPs at non-zero C values appear as multiple distinct “humps” (Figure 6A1) phase-locked to different instantaneous frequencies (Figure 6A2). These humps are consistent for different sweep rates and for up- and downward sweep directions, as illustrated in Figure 6C by superimposing the responses to C=1 with the time-reversed responses to C=-1. As in Figure 5D, we turned to an independent analysis of frequency tuning using pure and AM tones. Indeed, some cells show several maxima in their iso-intensity contours, further supporting the interpretation that in most octopus cells the inputs are not smoothly distributed across frequencies but derive from “hotspots”. An example is illustrated in Figure 6D-F with responses to Schroeder (Figure 6D) and AM stimuli (Figure 6E,F). Responses to AM tones reveal three discontinuous frequency regions that generate large EPSP integrals (at carrier frequencies near 3, 19, and 33 kHz: voltage integral in Figure 6E and raw traces in Figure 6F). Multiple “humps” were also observed in the Schroeder responses (Figure 6D), and these were triggered by instantaneous frequencies that closely match the frequency tuning observed with AM tones (matching symbols Figure 6D,E). In a minority of octopus cells (37.5%, n = 6/16, Figure 6B1), there are no obvious humps in the responses at any C value, and changes in sweep speed just change EPSP width (Figure 6B2). Our interpretation is that in these cells the inputs are derived from a smooth and continuous cochlear sector, while for most cells the inputs are clustered at discontinuous cochlear “hotspots”.

The finding of “humps” in membrane potential provides a straightforward interpretation of the multi-peaked cycle histograms often observed in the spike output of octopus cells (Figures 5,S4). Taken together with the observation that highly entrained action potentials can be triggered by a range of C-values and frequencies even within a neuron (Figure 2D,2F,5A,5C,S4), this indicates that compensation for delay followed by coincidence across input frequency channels is not a prerequisite for entrained firing and direction selectivity.

### Directional tuning to FM sweeps is dependent on the sequence of major versus minor inputs

We next studied how the “humps” in the responses to Schroeder stimuli, caused by the “hotspots” in frequency tuning of the inputs, generate tuning to frequency sweeps in the spike output. Six recordings with humps show clear spikes that are distinguishable from EPSPs (2 sharp, 4 whole-cell). Two examples are illustrated in Figure 7, where subthreshold traces (red) overlay suprathreshold (black) traces. Different humps (most easily discerned at |C|=1) are marked with letters. For the two cells illustrated, there is a major “hump” *a* and a smaller hump *b*. For each hump, the triggering frequency is consistent across sweep speeds and directions (magenta lines Figure 7A,B).

**Figure 7.**
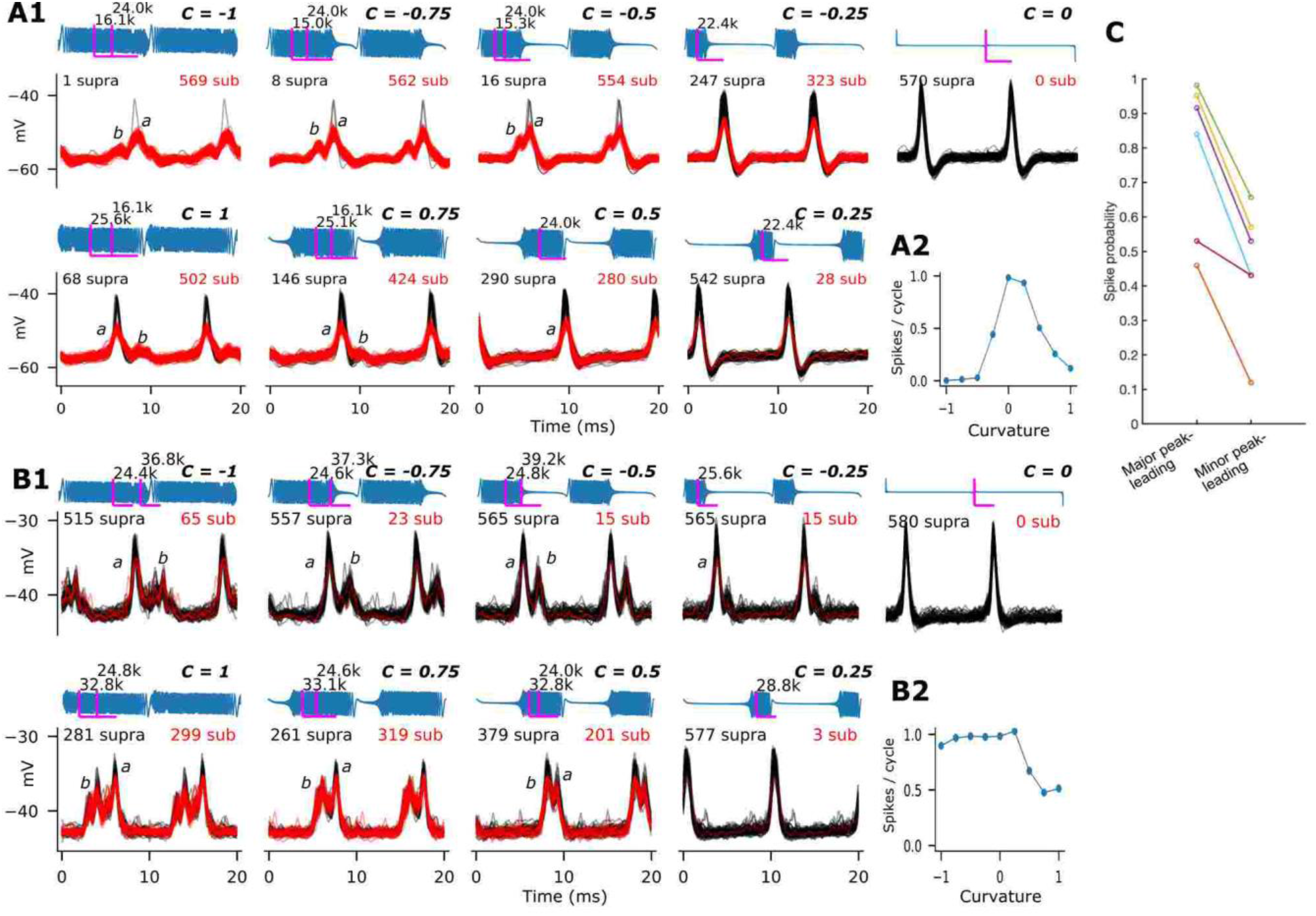
Intracellular responses show that octopus cell’s direction selectivity is dependent on the activation sequence of hotspot inputs. Overlay of sub- (red) vs supra- (black) threshold traces from two octopus cells to Schroeder stimuli (F0=100Hz, harmonics #1 - #400, 30 dB SPL). Each condition was tested 580 cycles. For clarity, a total of 100 traces are presented in each panel, with sub- and supra-threshold traces sampled randomly according to their rate of occurrences (numbers in each panel). Cells in (A) prefer downward sweeps (C > 0), whereas the cell in (B) prefers upward sweeps (C < 0). Responses of each cell show a major EPSP (“a”) and at least one minor EPSP (“b”), indicative of major vs minor inputs to the cell. Estimated triggering frequency for each input is marked on top of the magenta lines (cf. Figure 3) in each panel. **(A1)** The downward-preferring cell with a major input (“a”) at 22-26 kHz (∼ CF) and a minor input (“b”) at 15 - 16 kHz. **(A2)** Spike rate responses in (A1). **(B1)** The upward-preferring cell with a major input (“a”) at 24-25 kHz (∼ CF) and a minor input (“b”) at 33-39 kHz. Note that the major input triggers more spikes in upward sweeps (C < 0), and in some cases minor input can also trigger spikes (C = 1, -1, -0.5). **(B2)** Spike rate responses in (B1). **(C)** The spike probability is dependent on the activation sequence of the major vs minor inputs and is significantly higher when the major peak leads in the EPSP sequence (major-lead vs minor-lead: 0.79 ± 0.21 vs 0.47 ± 0.17, p < 0.005, paired t-test, n = 6 cells). The probability for each cell is pooled from responses to all curvatures (at least 580 repetitions/cycles per curvature) that fits the description in the x axis.

Given that octopus cell spiking requires a steep depolarization (Ferragamo and Oertel, 2002; McGinley and Oertel, 2006), a reasonable prediction is that major input *a* provides depolarization that is sufficient in amplitude and speed to drive spikes. Indeed, spikes are usually associated with major EPSP *a*. However, the spike probability of major EPSP *a* is not always the same (e.g. few spikes for C=-1 but many for C=1 in Figure 7A, and the opposite for Figure 7B). Moreover, sometimes the spikes can be evoked by both major and minor EPSPs (e.g. in Figure 7B1 by event *a* at C=-0.5 and *b* at C=0.5). This explains the difference in triggering frequency observed in the spiking data, mentioned above (Figures 5,S4). The critical observation in Figure 7, however, is that high spike probability, i.e. the preferred sweep direction, is associated with early triggering of major event *a* followed by later triggering of minor event *b*, while the reverse sequence results in low spike rates. For example, for the cell in Figure 7A, minor EPSP *b* precedes major EPSP *a* for C=-1, -0.75, and -0.5. and in these conditions the spike rate is low; but when *b* follows *a* (C=1, 0.75, 0.5), the spike rate is high. For the cell illustrated in Figure 7B the sequence is reversed: *a* leads *b* at C<0 but lags at C>1, and indeed the rate tuning of this cell (Figure 7B2) is opposite to that of Figure 7A.

These observations suggest that directional preference is not determined by coincidence of inputs with different CF, but by the ability of major inputs to generate a spike, only if not immediately preceded by minor inputs. Thus, it is the sequence of events, rather than their coincidence, that underlies tuning to frequency sweeps. Because major and minor inputs reverse their sequence as the sweep direction changes, we hypothesize that the activation sequence of EPSPs interacts with the abundant KL channels (Cao and Oertel, 2005; Golding et al., 1999; Oertel et al., 2008) to determine the spike-triggering efficacy of the major EPSP. Indeed, in the 6 cells whose spikes are clearly distinguishable and exhibit clear EPSP humps, the spike probability is significantly higher when major input *a* leads compared with the condition where minor input *b* leads (0.79 ± 0.21 vs 0.47 ± 0.17, p < 0.005, paired t-test) (Figure 7C). Hence, direction selectivity reflects an interaction between the activation sequence of hotspots of inputs and intrinsic membrane conductances.

In a minority of cells there are no obvious major and minor inputs evoked by Schroeder stimuli (Figure 6B). An example with distinguishable spike is shown in Figure S5A, where the frequency sweeps evoke a single wide EPSP at long sweep durations. At high levels (e.g. 50 dB SPL, Figure S5C), the EPSP is less smooth and shows some evidence of separate events but not as clearly as in Figure 7. This corresponds well with the voltage responses evoked by long tone stimuli (Figure S5B), which reveal one major depolarization at around 19–33 kHz, without “hotspots”. This cell showed no directional preference (see spike counts in Figure S5A). Thus, limited direction selectivity (Figure 2K,L) may reflect more equally or smoothly distributed inputs across frequencies.

### Directional tuning can be reproduced with a biophysical model having only a single compartment with hotspot inputs

The dependence of directional tuning on the activation sequence of major vs minor inputs is likely due to the abundant KL channels present in octopus cells. When the sequence of activated inputs is such that minor inputs are activated first, they may not immediately evoke spikes but activate KL, thereby shunting the overall membrane conductance and preventing subsequent depolarization from generating spikes. We used a published, conductance-based single compartmental model of octopus cells (Manis and Campagnola, 2018) to test our interpretation. We provide ANF inputs (from our *in vivo* recordings, see Methods) centered at four different CFs (25, 20, 16 and 12 kHz) and tested the effect of different input weights. When most inputs are tuned at 25 kHz and a smaller number to the other frequencies, the model output is very asymmetrical with poor spiking when the “minor” inputs lead the “major” input in time (C<0), and high spiking for the reverse situation (C>0) (Figure 8A). When the input distribution is reweighted so that the number of inputs is largest at 16 kHz (Figure 8B), the model output now favors C=-0.5. Consistent with the *in vivo* observations, the preferred direction is again the one in which the major inputs occur first (C<0). When all inputs are equally weighted (Figure 8C), spiking is only observed at C=0. Taken together, this simple computational model, taken from the literature “as is” and fed with real ANF spike trains, reproduces our *in vivo* observation and supports our interpretation that asymmetry in the number of excitatory inputs clustered at different CFs together with intrinsic membrane conductances result in direction selectivity to frequency sweeps.

**Figure 8.**
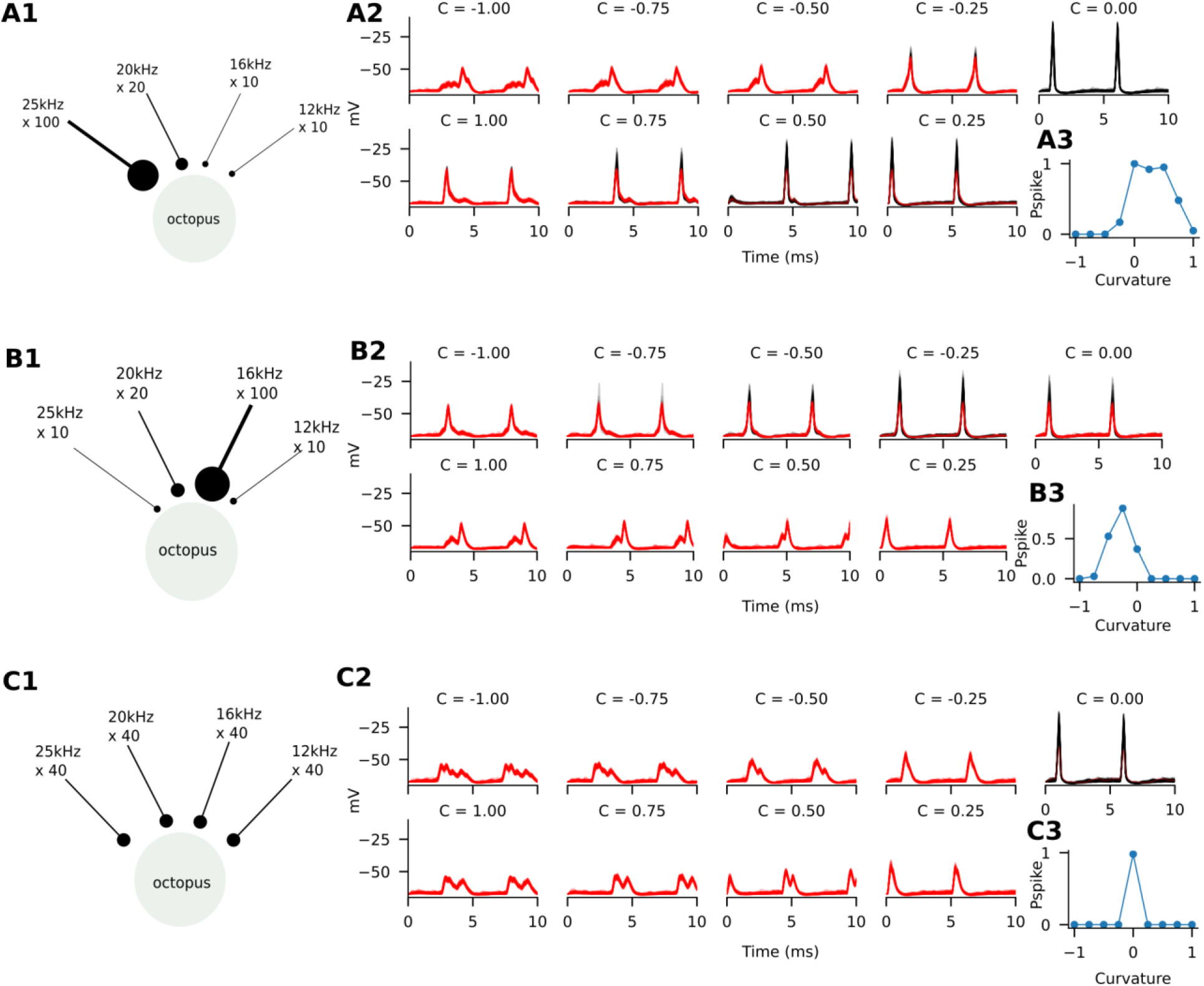
Direction selectivity can be recreated in a single-compartmental model of octopus cell with inputs tuned to “hotspot” frequencies. Spike times are randomly chosen from the response cycles of ANFs tuned near 25, 20, 16, and 12 kHz. The number of inputs for each CF is indicated in the cartoons (A1, B1, C1). The ANF responses were from in vivo recordings evoked by Schroeder stimuli with F0=100Hz, harmonics #1 - #400 at 30 dB SPL. Voltage responses of 100 trials from the model octopus cell for each condition are shown (A2-C2). Red traces indicate subthreshold responses; black traces indicate suprathreshold responses. The resulting spike rate tuning is shown in A3-C3. Even though each cell receives inputs tuned to the same CFs, direction selectivity is observed only in cells (A, B) with unequally distributed inputs. With equally distributed inputs, cell in (C) spikes only at C=0. Furthermore, the selectivity correlates well with the activation sequence of major vs minor inputs. When major inputs lead (C > 0 for cell A, C < 0 for cell B) the cell spikes. When minor inputs lead (C < 0 for cell A, C > 0 for B), the subsequent major EPSPs cannot trigger spike. This confirms that the intrinsic membrane conductances interact with input sequences to generate direction selectivity.

## Discussion

Octopus neurons have been portrayed as detectors of spike coincidences among nerve fibers derived from a wide cochlear sector. The two key observations reported here are that these cells show marked direction selectivity to frequency sweeps, and that this selectivity is based on a purely excitatory mechanism not of coincidence detection but of sequence detection. These findings considerably broaden the possible functional role of these cells and reveal a new type of neural computation that creates acute sensitivity to temporal patterns across inputs.

### Direction selectivity to FM sweeps

Neural sensitivity to “motion” of stimuli across receptor surfaces is a well-studied subject, particularly in visual systems (Mauss et al., 2017; Yonehara et al., 2013). The lack of a simple correspondence between external space and cochlear space fundamentally affects the nature of “motion” in hearing. Evidence for sensitivity to motion of sounds in external space is limited and controversial (Joris, 2019). With the cochlea being mapped in frequency, the closest equivalent to motion on the retina or skin is “cochlear motion” caused by changes in sound frequency.

Use of various forms of FM has uncovered direction selectivity at multiple anatomical levels, including the cochlear nucleus of diverse species [bat (Suga, 1964), guinea pig (Paraouty et al., 2018), rat (Møller, 1974), and cat (Britt and Starr, 1976; Erulkar et al., 1968; Godfrey et al., 1975; Møller, 1974; Recio-Spinoso and Rhode, 2020; Rhode and Smith, 1986)]. The speeds tested were several orders of magnitude slower than those used here (>1 kHz/ms), and revealed modest direction selectivity in the small number of presumed octopus cells sampled, mostly favoring upward sweeps (Godfrey et al., 1975; Paraouty et al., 2018; Recio-Spinoso and Rhode, 2020; Rhode and Smith, 1986). One study in chinchilla found “little additional processing” in cochlear nucleus responses to Schroeder stimuli but did not report responses from octopus cells (Recio, 2001).

We tested octopus cells with FM sweeps because of earlier speculations of FM selectivity on structural grounds (orientation of dendrites relative to ANFs) (Szentagothai and Arbib, 1974) as well as the more specific hypothesis that octopus dendrites counteract the dispersive effects of cochlear delay (McGinley et al., 2012). We discovered remarkable sensitivity of octopus cells to the rate and direction of linear FM sweeps (Figure 2). The nature of this sensitivity and the underlying mechanisms are not consistent with the latter hypothesis: octopus cells did not only entrain to click trains (Figure 1), but also to stimuli that generate large delays across the ANF population (Figures 3,4). The cells are thus not only triggered by the temporal pattern of cochlear delays in response to clicks, but also by patterns that strongly deviate from the latter. Moreover, neurons differed in their preference with some preferring downward while others upward frequency sweeps. These results not only challenge the hypothesis of cochlear delay compensation, but more generally the proposal that these cells detect specific patterns of coincidences across ANFs. Indeed, while cells generally preferred one sweep direction, there was large tolerance for sweep speed. Because the temporal delays across ANFs are much larger for slow sweeps than for clicks (Figure 4), the observation that very different temporal patterns could trigger almost identical, entraining responses, suggests that another mechanism than coincidence detection underlies direction selectivity.

### Sequence detection

Previous studies of mechanisms generating low-speed auditory FM selectivity, from midbrain to cortex, pointed to an interaction between excitatory and inhibitory synaptic inputs (Gittelman et al., 2009; Kuo and Wu, 2012; Zhang et al., 2003): early activation of inhibitory inputs prevents subsequent excitatory inputs from generating spikes, leading to an asymmetry in spike rate to opposing sweep directions. However, despite some morphological evidence for GABAergic and/or glycinergic terminals in the octopus cell area (Adams and Mugnaini, 1987; Alibardi, 2003; Juiz et al., 1996; Saint Marie et al., 1989) [but see (Wickesberg et al., 1991) and (Ngodup et al., 2020)], there is no physiological evidence for evoked inhibition in octopus cells *in vitro* (Golding et al., 1995). We did not observe apparent evoked hyperpolarization *in vivo* either. Rather, our data and model show that direction selectivity in octopus cells reflects an interaction of the sequence of excitatory inputs with intrinsic membrane conductances.

Both spike (Figures 5,S4) and membrane responses (Figure 6,7) to FM sweeps, as well as measures of frequency tuning (Figures 5,6), suggest the existence of discrete sets of inputs tuned to different frequencies to most octopus cells. Moreover, these inputs differ in effective strength as reflected in amplitude of depolarization and ability to trigger spikes (Figure 6A,7). The key mechanism generating direction selectivity is the temporal sequence in which strong and weak inputs are activated: leading activation of weak inputs prevents subsequent strong inputs from triggering spikes. In some neurons, discrete sets of inputs differing in strength were not obvious (Figure 6B,S5), and this was associated with weak direction selectivity. The cellular mechanism suppressing spiking after activation of a weak input is not synaptic inhibition but likely the activation of KL channels, specifically Kv1, known to be abundantly present in octopus cells (Bal and Oertel, 2001; Oertel et al., 2008) and cause the cells to fire more readily to depolarizing current ramps with steep but not shallow slope *in vitro* (Ferragamo and Oertel, 2002; McGinley and Oertel, 2006). Interestingly, a similar sequence sensitivity was observed in neurons of the medial superior olive (MSO), which also has abundant Kv1 channels (Franken et al., 2015).

Our mechanistic interpretation was supported by feeding a simple published model of octopus cells (Manis and Campagnola, 2018) with our experimentally recorded ANF spike trains (Figures 2,4), from which we took discrete ANF populations clustered around a small number of CFs. When inputs tuned to one frequency dominated over inputs tuned to other frequencies, direction selectivity emerged and its sign was consistent with the sequence in which these inputs were recruited relative to other inputs (Figure 8A,B). In contrast, when inputs were equal in number, firing was maximal for click-like stimuli (Figure 8C), even though no compensation for cochlear delay was applied.

The structural correlate to “hotspots” remains to be determined. EM studies of axons traversing the octopus cell area show both large and small boutons innervating somatodendritic compartments (Kane, 1973) and single labeled ANFs form either large “modified endbulb” endings on octopus somata, or “streaming collaterals” where many small terminals emerge from collaterals that course along octopus dendrites (Fekete et al., 1984; Rouiller et al., 1986). The difference in effective strength of inputs tuned to different CF likely reflects the number, size and placement of ANF terminals. The physiological findings (low variability in subthreshold responses, Figures 7,S5, and entrainment in spike output, Figures 2,5) suggest that each “hotspot” reflects activation of many inputs of similar CF.

We do not imply that coincidences among inputs are irrelevant. The octopus cell’s combination of low input resistance with many small, subthreshold inputs indeed necessitates the simultaneous or coincident activation of many inputs to elicit spiking. However, mere consideration of coincidences is insufficient to account for the direction selectivity to FM sweeps. Across-frequency coincidences are not essential since inputs from a single “hotspot” can be sufficient to trigger spikes (Figure 7), except when preceded by an input which does not itself elicit spikes. Thus, even while considering that each “hotspot” reflects coincident activation of many inputs, it is “hotspot” magnitude and sequence of activation that holds the key to direction selectivity. Only at low stimulus level, when there are likely too few spikes generated within “hotspots”, does alignment across CFs (click trains, C=0, e.g. Figure 2F) become a necessary condition for spiking. It is a moot point whether there is a compensation in that condition for small high-frequency cochlear delays, since such compensation would only be relevant under threshold conditions.

### Functional implications

The frequency sweep rates employed here are very fast (mostly 1 to 20 kHz/ms). Natural sounds, including human and animal vocalizations, are dominated by much lower sweep rates (Varnet et al., 2017). However, there is a methodological caveat: the sweeps in instantaneous frequency as used here are not reflected in the long-term spectra, which consist of a stable harmonic series (Figure S1B). The F0s we used (50 – 400 Hz) span a range that humans (and animals) are commonly exposed to, and it is likely that in natural conditions the phases of harmonics over a certain frequency range sometimes combine to produce frequency sweeps (“chirps”). Thus, fast sweeps in instantaneous frequency are likely more common than suggested by long-term spectra.

We propose that the sensitivity uncovered here is of wider significance than detection of frequency sweeps. Fundamentally, we find that octopus cells are sensitive to timing relationships between ANFs of widely different CF, and that the preferred timings and frequencies differ between cells. We surmise that this sensitivity provides a basic “display” mechanism which can be recruited for a variety of perceptual tasks, which likely differ between species. For example, temporal interactions between harmonics of different sound sources, e.g. different speakers, have repeatedly been proposed as a mechanism to segregate such sounds (Carney, 2018; de Cheveigné, 2021; Larsen et al., 2008; Palmer, 1990; Sinex, 2005, 2008; Sinex and Li, 2007). Our findings suggest that octopus cells are very sensitive to such interactions.

A conundrum that has plagued temporal models of hearing based on coincidence detection is the requirement for internal delays to bring spike trains into coincidence (de Cheveigne and Pressnitzer, 2006). Even in the binaural system, there is much controversy regarding the source of such compensating or internal delays despite extensive study (Grothe et al., 2010; Joris and van der Heijden, 2019; Karino et al., 2011). Our findings in octopus cells reveal a new mechanism that creates sensitivity to temporal relationships between inputs without recourse to explicit delay compensation.

## Acknowledgments

This work is supported by US National Institute of Health (NIH) grant DC006212 (P.H.P and P.X.J), NIH BRAIN grant 1R01NS118402 (P.X.J.), Research Foundation Flanders (FWO) grant G085421N (P.X.J), and EMBO long-term postdoctoral fellowship ALTF 7-2017 (H.-W.L.). The authors thank members of the Joris lab for their assistance.

## Author contributions

H.-W.L., P.H.P., and P.X.J. conducted the experiments; H.-W.L. analyzed the data; H.-W.L. and P.X.J designed the experiments; H.-W.L., P.H.P, and P.X.J. wrote the paper.

## STAR Methods

### Animals

Seventy adult Mongolian gerbils of either sex (aged between 40 days to one year) were used for this study. All animal handling and surgical procedures were approved by KU Leuven’s Institutional Animal Care and Use Committee.

### Surgery

Animals were anesthetized throughout the experiment and placed in a double-walled sound- proof booth (IAC, Niederkrüchten, Germany). Before surgery, the anesthesia was induced by intraperitoneal (i.p.) injection of Ketamine/xylazine mixtures dissolved in saline (ketamine 80 mg/kg, xalyzine 12 mg/kg). During the experiment, anesthesia was maintained by intramuscular or i.p. injection of ketamine (25 mg/kg) and diazepam (Valium, 1.0mg/kg). Anesthetic depth was regularly monitored by pedal reflex. Body temperature was maintained at 37 ℃ with a heating pad. A metal headbar was attached to the frontal skull by dental cement. We used a dorsal approach to expose the cochlear nucleus. A 4-mm diameter craniotomy was made on the lateral part of the occipital skull. The cerebellum dorsal to the cochlear nucleus was then carefully aspirated. Silicon oil was applied to the craniotomy during sharp electrode recordings to reduce brain pulsation. Some ANF recordings were obtained via the round window niche approach developed by Sokolich (Chamberlain, 1977; Zwislocki and Sokolich, 1973).

### Electrophysiology

Extracellular single unit recordings were made with sharp electrodes (tip resistance 80 - 120 MΩ). Intracellular recordings were made either with sharp electrodes or patch clamp pipettes (tip resistance 6 - 7 MΩ). All electrodes were pulled from borosilicate micropipettes (WPI 1B100F-6 for sharp, WPI 1B120F-4 for patch electrodes) on a horizontal puller (Sutter Instrument, P-2000 or P-87). The internal solution for sharp electrodes was 0.5 - 1M KCl, with 1.5 - 2 % neurobiotin when we aimed for morphological labeling. Internal solution for patch clamp recordings consisted of (in mM): 115 K gluconate, 4.42 KCl, 0.5 EGTA, 4 Mg-ATP, 0.3 Na-GTP, and 0.1-0.4% (w/v) biocytin, with pH adjusted to ∼7.30 by KOH and osmolality to ∼290 mOsm by sucrose. Electrodes were mounted on a hydraulic microdrive supported by a micromanipulator, and advanced into the dorsal cochlear nucleus under visual guidance. For sharp electrodes we advanced the electrode in 1-2 µm steps while recording from any cells encountered until reaching ∼1000 µm below the surface. The pipette was then withdrawn and replaced, and another penetration initiated. For patch pipettes, positive pressure (4-5 psi) was applied before entering brain tissue. We advanced the electrode at ∼100 µm / second until reaching the depth at around 500 µm below the surface. Pressure was then reduced to ∼ 0.4 psi and the electrode advanced 1-2 µm per step until signs of touching a cell were observed (tip resistance fluctuating at the rate of heart pulsation). The positive pressure was then immediately released to form a gigaohm seal and brief suction applied to enter the whole-cell configuration (Franken et al., 2015; Lu and Joris, 2021; Margrie et al., 2002). Bridge resistance was then balanced and pipette capacitance compensated. Initial bridge resistance was usually around 30 - 50 MΩ but often fluctuated during the recording due to pulsations caused by breathing or heartbeat. Recordings were discontinued if the bridge resistance went above 100 MΩ. Axoclamp 2B (Axon Instruments) and Dagan BVC- 700A amplifiers (Dagan) were used for amplifying signals from sharp and patch electrodes, respectively. Voltage responses were sampled at 100 kHz online, decimated to 20 kHz and low-pass filtered at 5kHz for analysis. For patch clamp recordings, voltages were corrected for an estimated junction potential of 10 mV. Spikes were detected online using a custom-made peak- picker and digitized at 1 µs resolution. Detailed methods are provided in Lu and Joris (2021).

#### Sound stimuli

Sounds were digitally synthesized in Matlab and then delivered to an acoustic driver (Supertweeter or Horntweeter, Radio Shack) using Tucker-Davis Technologies System 3 hardware. The speaker was connected to the animal’s ear canal through a custom-made earpiece which incorporates a probe tube near the eardrum. Before recording, amplitude and phase of the transfer function of the closed acoustic system were measured with a 0.5-inch condenser microphone (Bruel and Kjaer) coupled to the probe tube. All sound stimuli were compensated for using this transfer function.

The main stimuli of interest were Schroeder harmonic complexes, click trains, and various types of tones, as described in the main text, but for each neuron a brief characterization of the frequency tuning was first obtained. We used either fixed SPL tones stepped logarithmically in frequency, which provided the frequency of maximal spike rate response (best frequency, BF), or a threshold frequency tuning curve using a tracking algorithm (Geisler and Sinex, 1982), which provided frequency of lowest threshold (characteristic frequency, CF). Because octopus cells tend to be poorly driven by pure tones (Figure 1A), this was often supplemented with amplitude-modulated tones varied over a wide range of carrier frequencies (see below, AM tones). When action potentials were clearly discriminable, these tuning measures were based on triggered spikes. If, during intracellular recordings, spikes were too small to be discriminated from EPSPs, triggering included both supra- and subthreshold events.

#### Schroeder stimuli

Schroeder harmonic stimuli with fundamental frequency F0 of a particular phase curvature C were generated according to following equation:

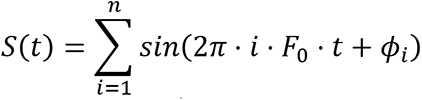

where t stands for time, and the phase of the i^th^ harmonic, *ɸ*_i_, is designated by

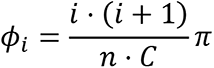

and

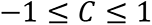

Figure S1A shows representative waveforms at various curvatures. Given the harmonic spectrum (Figure S1B) the waveforms are periodic with fundamental period 1 / F0. Two periods are shown for all C values (blue), with the instantaneous frequency superimposed (orange). At C = 0, the waveform is similar to a click train repeating at F0. At non-zero C values, the waveform is essentially a series of linear frequency sweeps repeating at the rate of F0, and the sign of C determines the direction of the sweep (upward sweep for C < 0; downward sweep for C > 0). In this study we mostly used F0 at 100 Hz with 400 harmonics (i = 1 to 400), which gives a spectral range from 100 Hz to 40 kHz. Occasionally, other F0s ranging between 50 to 400 Hz were used, but the spectral ranges were always restricted within F0 and 40 kHz. Care was taken that the stimulus spectrum covered the frequency range to which the recorded cell responded (as assessed by pure or amplitude-modulated tones), within the usable range of the driver (as assessed with the acoustic calibration). The combination of F0, frequency range, and C values determines the sweep rate of the instantaneous frequency. For the largest C values (|C| = 1), a frequency sweep between lowest and highest harmonic frequency spans the entire fundamental period; for |C| = 0.5 the duty cycle is halved and the sweep rate doubled; and for C = 0 the sweep rate is nominally infinite. Sweep rate and direction are provided in Results and Figures. Schroeder stimuli were presented with C varying from -1 to +1 in steps of 0.25 and repeated at 1000 ms intervals (= sound duration 600ms + silent interstimulus period 400 ms), and the sequence was repeated at least 10 times. Initial sound level was usually near 30 dB SPL but was varied if time allowed. Sound levels are specified as the level of individual harmonics, hence a complex of 400 harmonics with each harmonic at 30 dB SPL has an overall level at 56 dB SPL.

#### Click trains

We used trains of 20 µs rarefaction clicks. Each round of stimuli started with click trains varying from 100 to 1000 Hz in steps of 100 Hz, using the same timing parameters as the Schroeder stimuli (600 ms duration, repeated every 1000 ms). If time allowed, each round was repeated 10 times. The sound level for click train is computed as the RMS level of the calibrated waveform. We usually started with 70 dB SPL and varied in +10 or -10 dB steps if time allowed.

#### Short-tone pips at BF or CF (STCF)

STCF consist of a 25-ms duration tone at the cell’s BF or CF with 2.5 ms rise/fall time. To characterize the response type of the recorded cell, STCF were presented 200 times at 100-ms interval for a given sound level, and the sound level was sequentially varied from around 90 dB SPL to the threshold level in 10 dB steps, from which peristimulus time histograms (PSTHs) were generated and displayed on-line.

#### Long tones and AM tones

Long, 300-600 ms tones were delivered at various carrier frequencies, mostly ranging from 0.5 - 40 kHz in 0.4 octave steps. Because sinusoidally amplitude modulated (AM) tones are a more effective stimulus to drive octopus cells, often at lower threshold, than pure tones (Recio-Spinoso and Rhode, 2020; Rhode, 1994), while still being narrowband, we sometimes resorted to the use of these long AM tones (modulation depth 100%, modulation frequency of 100 Hz, rise/fall time 5 ms) to study the frequency tuning of the cell and obtain a BF. The AM tones were generated according to:

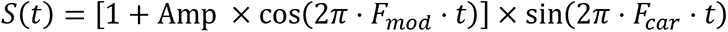

where

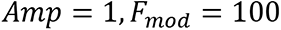

*t* stands for time in seconds; *Fcar* for carrier frequency in Hz.

For analysis in Figure 6F, the first 15 and last 5 ms of the responses during long AM stimuli were omitted to generate cycle-averaged waveform.

### Data analysis

Analyses were done in Matlab (Mathworks) and Python. Unless otherwise specified, data were presented as mean ± S.D.

Response type definition: For spike responses obtained by sharp electrodes, we follow the standard convention to categorize by PSTH to STCF (Godfrey et al., 1975; Lu et al., 2018; Rhode and Smith, 1986). Three cell types are discussed in this paper: ideal onset with little to no sustained activity (Oi), onset with low sustained activity (OL), and primary-like (PL). We define the onset units by onset-to-steady-state spike ratio > 10 (Rhode and Smith, 1986; Winter and Palmer, 1995). Here, onset spike rate is the spike rate within the 1-ms window centered at the peak of the PSTH. Steady-state spike rate is the spike rate between 10 ms post-onset and the end of the stimuli. Octopus cells are associated with Oi or OL responses (Godfrey et al., 1975; Pfeiffer, 1966; Rhode et al., 1983; Smith et al., 2005). Various criteria have been used to separate Oi and OL responses: we used the criterion of Rhode and Smith (1986) and defined Oi based on sustained rate < 30 spikes/s, which segregated the two groups along a number of properties (see Results, Figure 1B). We occasionally encountered onset with chopping (Oc) units, which are associated with the glycinergic D-stellate cells but not octopus cells (Smith and Rhode, 1989; Smith et al., 2005) and are not further considered in this study. PL response types in this study were either obtained at the depth of the octopus cell area or in the nerve bundle close to the round window, and hence are labeled as the ANF response in this study. We use spontaneous firing rate of 18 spikes / sec as a threshold to categorize low vs high spontaneous rate fibers (Huet et al., 2016; Liberman, 1978).

Contrast ratio calculation: we used a conventional contrast ratio, also termed direction selectivity index, to measure the degree of direction selectivity in response to the Schroeder stimuli (Britt and Starr, 1976; Heil et al., 1992; Nelken and Versnel, 2000; Paraouty et al., 2018). It is calculated as:

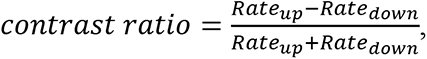

where Rateup, Ratedown are the spike rate measured in response to upward (C < 0) and downward (C > 0) sweeps, respectively. Hence, a positive contrast indicates a preference for upward sweeps, and negative for downward sweeps. A contrast at 0 indicates no preference for either direction. Because such ratio is sensitive to small denominators, only responses with more than 10 spikes / sec were included in this analysis.

For temporal analyses of spike patterns, we used two metrics. Vector strength (Goldberg and Brown, 1969) was calculated at the frequency of interest (pure tone frequency or F0 for responses to Schroeder stimuli). Only trials with > 10 evoked spikes were included. Significance was assessed with the Rayleigh test (Mardia, 1972). Shuffled autocorrelograms (SACs) were calculated with 50 μs binwidths (Joris, 2003). The correlation index CI (Joris et al., 2006) was obtained as the peak value of the normalized SAC at 0 delay.

For cycle-based histograms (Figure 3, 4, S4), the dotrasters illustrate spiking over two stimulus cycles for each C condition. The bottom rows of dots are for the initial stimulus cycle and the top tows for the last stimulus cycles within the 600 ms stimulus. The bottom traces show corresponding cycle histograms, i.e. spike counts over 2 cycles. Binsize = 0.1 ms.

### Modeling

We adopted a published octopus cell model for simulations in Figure 8 (Manis and Campagnola, 2018). The model was run in the NEURON 8.0 environment in Python (https://www.neuron.yale.edu/neuron/) (Carnevale and Hines, 2006), with the default parameters (RM03, II-o) from Manis and Campagnola (2018), i.e. it contains HCN, KL, KH (high-voltage activated potassium channels), leak, and voltage-gated sodium channels (jsrna in the model). It is a single-compartmental model, with only soma present, and the spike trains recorded from the ANFs in this study were fed to an alpha synapse (ExpSyn) connected to the center of the soma (position = 0.5) via the VecStim mechanism in NEURON. The tau for the alpha synapse is set to 0.35 ms, which matches the EPSC decay measured in mouse octopus cells in the brain slice (Gardner et al., 1999). The conductance of the alpha synapse is set to 2 nS, which is the upper limit of the estimated unitary synaptic conductance measured in the mouse brain slice (Cao and Oertel, 2010). Temperature is set to 33°);C. For simulating Schroeder evoked responses, spike trains were randomly sampled with replacement from the recorded responses in a given ANF in this study on a cycle basis. Only ANF responses with 10 repetitions (stimulus duration for each repetition: 600ms) to a given Schroeder stimulus condition (C = -1 to 1, F0=100Hz, 30 dB SPL) are selected. The first 10 ms response is omitted to avoid onset effects, hence there are a total of 590 available cycles to sample from a given ANF recording. The same ANF response is sampled repeatedly to mimic multiple inputs tuned to the same CF. All spike trains were then fed to the point alpha synapse connected to the soma. A voltage threshold is set to -40mV to detect spikes.

## Supplementary figures

**Figure S1.**
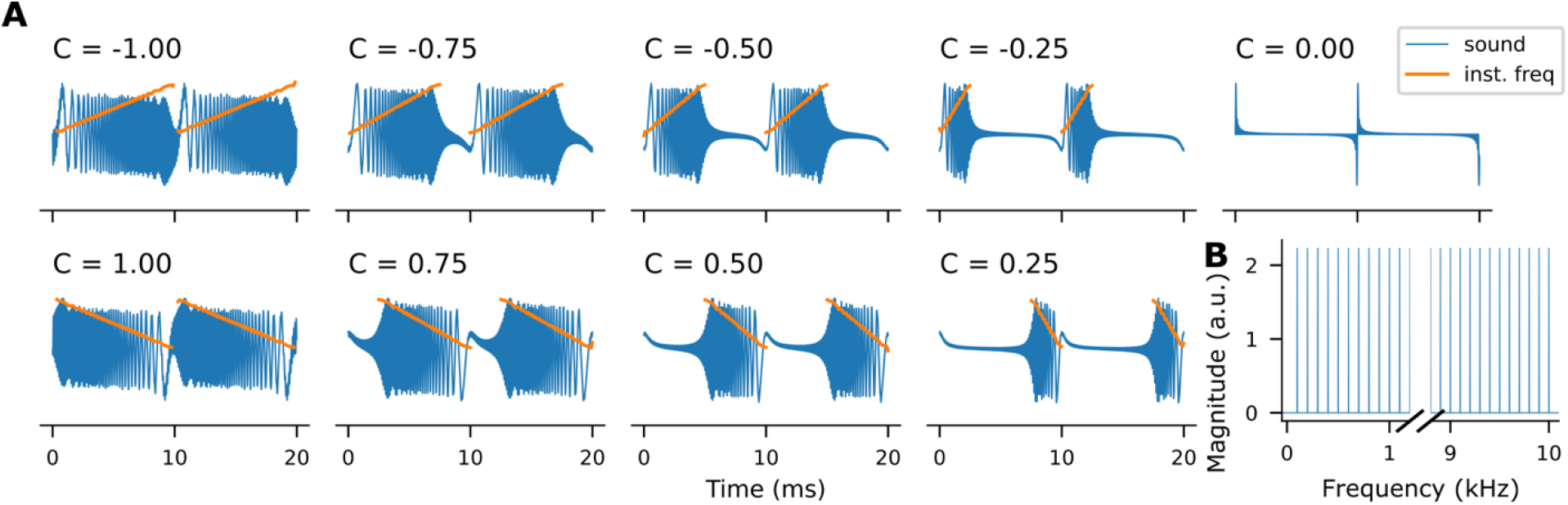
Schroeder stimuli with F0 = 100 Hz and harmonic number 1 - 100 at different phase curvatures. **(A)** Schroeder stimuli with negative curvature (C < 0) are a series of linear upward FM sweeps repeating at F0 (top row, each panel shows 2 cycles); positive curvature (C > 0) results in downward FM sweeps. As C approaches 0, the duration of FM sweep shortens and the sweep speed increases. At C = 0 the waveform becomes a click train. The ordinate scale is normalized to the maximum of the stimulus waveforms. Blue: sound waveforms. Orange: instantaneous frequency during the sweep derived by the Hilbert transform. **(B)** Independent of the C value, the spectrum of Schroeder stimuli consists of a simple harmonic series of equal magnitude.

**Figure S2.**
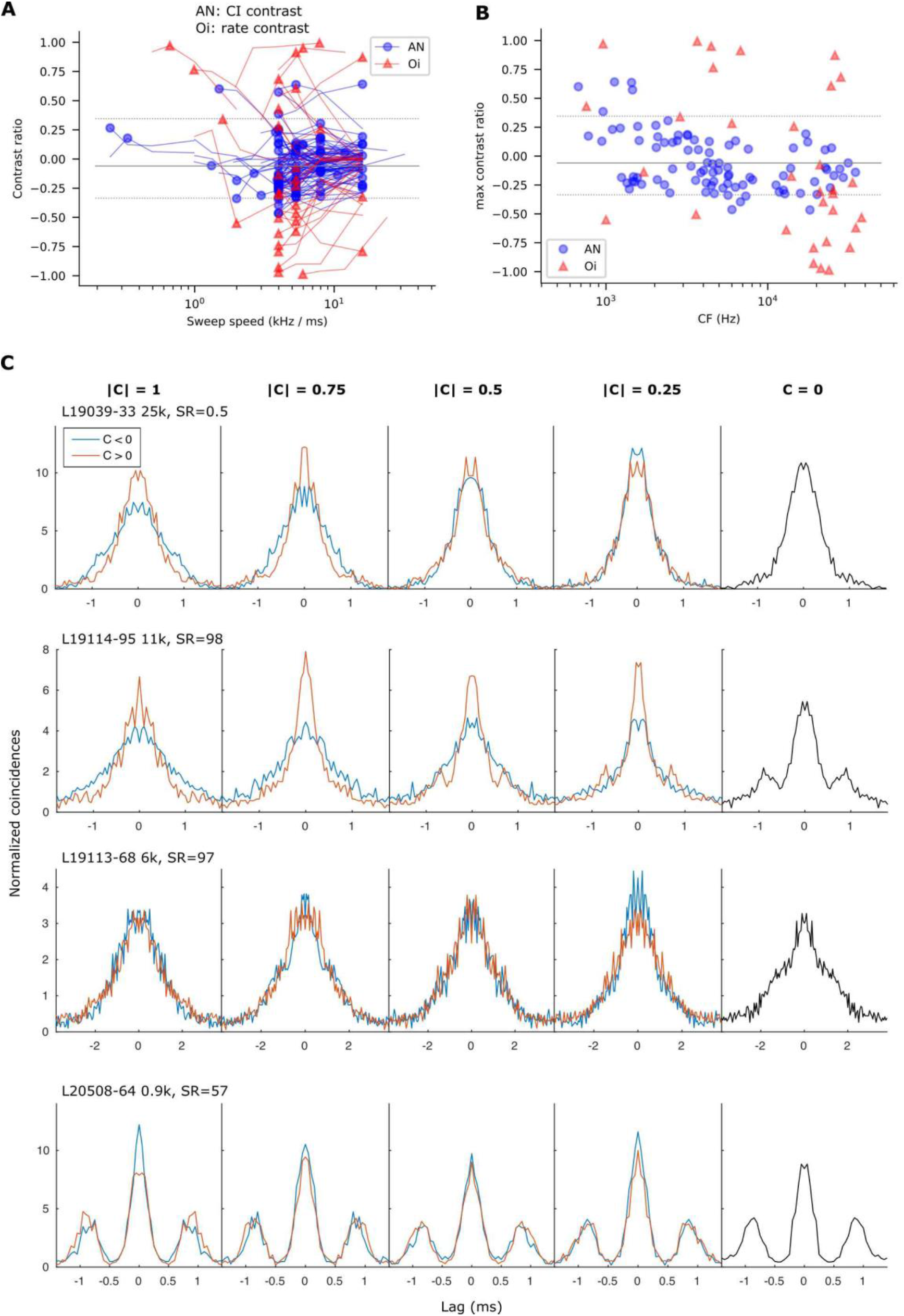
Coincidence analysis of ANF responses to Schroeder stimuli. **(A and B)** Population plots of contrast ratio in octopus cells (red) and auditory nerve (AN, blue) vs. sweep speed (A) and CF (B), using same format as in Figure 2K,L. Octopus data are same as in that figure. For the AN, the contrast ratio is calculated from the correlation index CI (peak at zero delay in SACs as in panel C). **(C)** Example shuffled autocorrelograms (SACs) from four ANFs arranged in CF from high (top) to low (bottom). Each panel shows a pair of SACs to C values of opposite sign (C < 0: blue, C> 0: red), from which two CI values are obtained to calculate a CI contrast ratio. The panels in the rightmost column (black) show the single SAC for C = 0. Labels on the top left of each panel indicates the dataset number, CF, and spontaneous firing rate (SR, spikes / sec) for the ANF.

**Figure S3.**
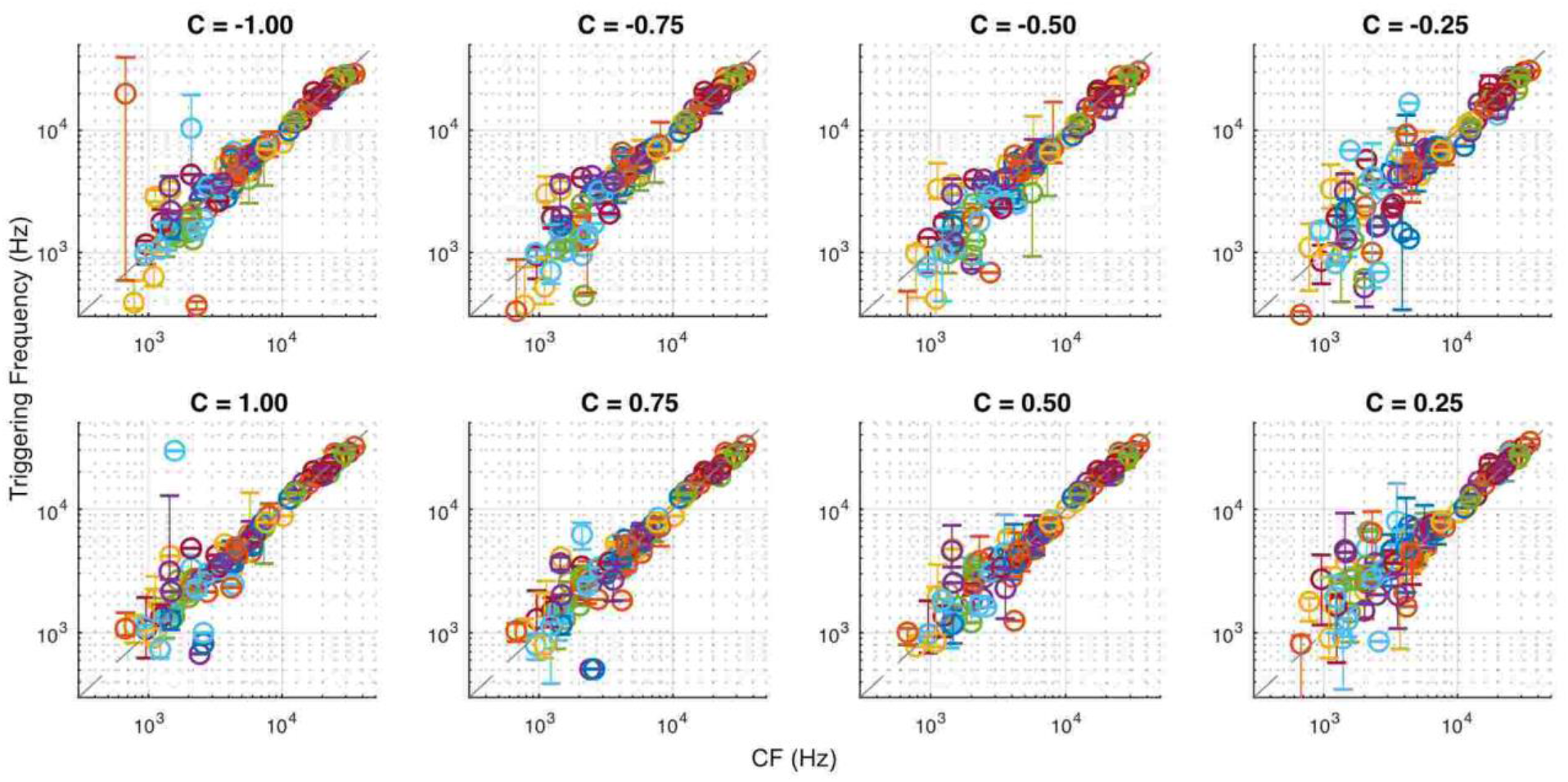
Schroeder triggering frequencies match the CFs of the ANF across all phase curvatures. Same format as Figure 3B.

**Figure S4.**
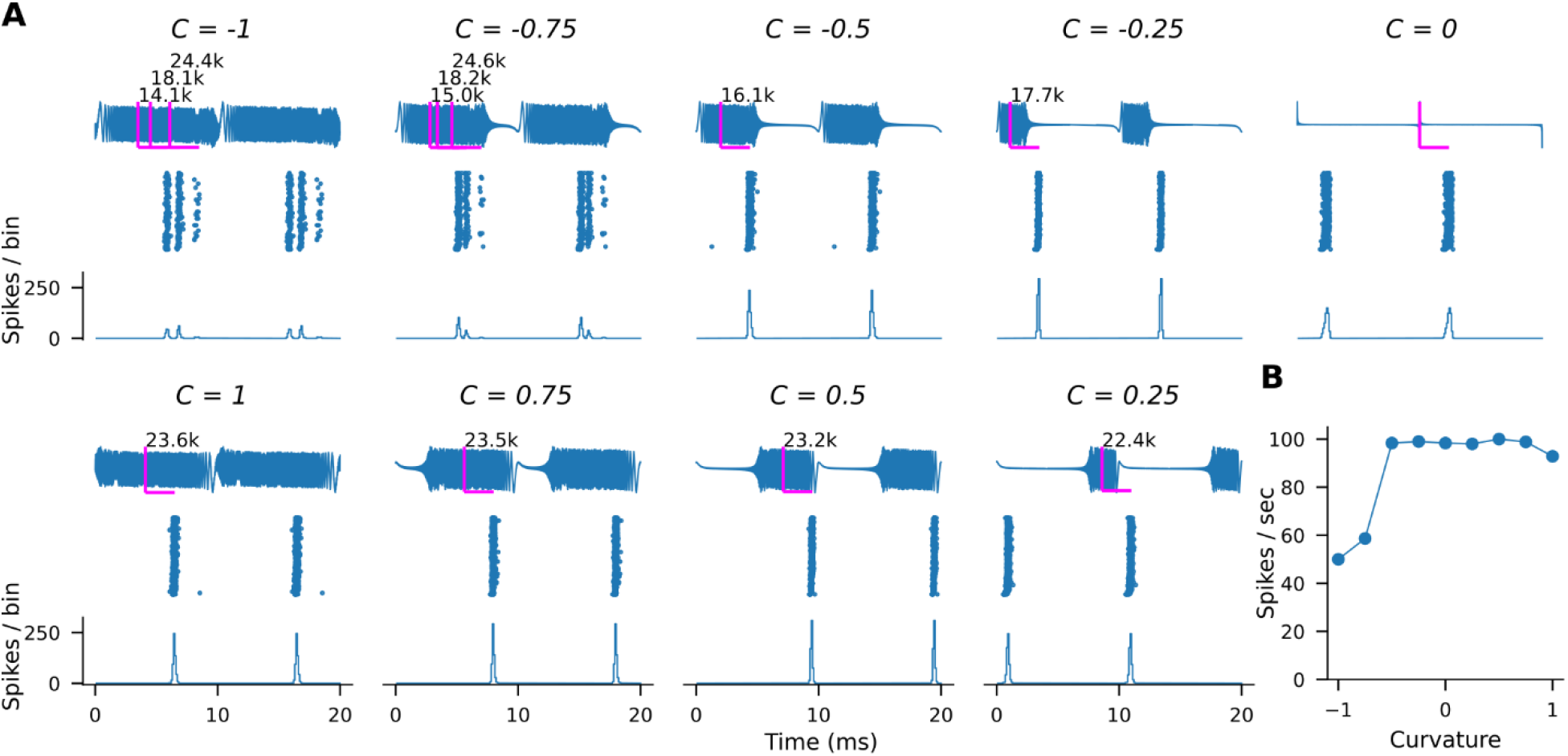
Octopus cell with three spike-triggering frequencies in its Schroeder responses. **(A)** An octopus cell (CF: 22 kHz) with discontinuous hotspots in inputs at non-preferred direction (C < 0). Same layout as in Figure 5 (A): first row sound waveform, second row cycle dotraster, third row cycle histogram (2 cycles shown). **(B)** Rate vs Curvature plot of the response in (A).

**Figure S5.**
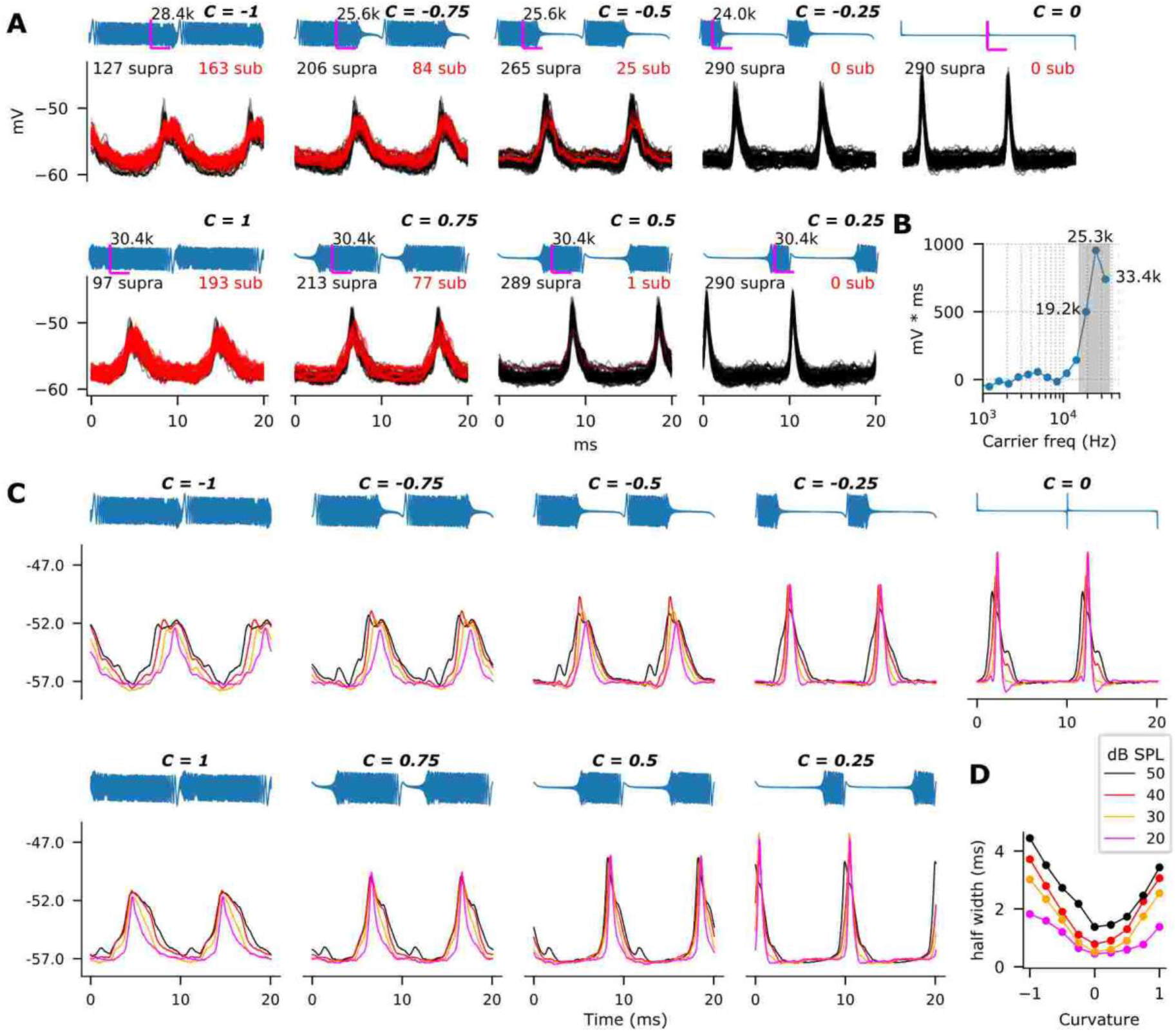
An octopus cell lacking direction selectivity, “hotspots”, and “humps”. **(A)** Intracellular traces (same format as Figure 7) for octopus cell lacking direction selectivity: spike counts (“supra”, black) are similar for C > 0 and C < 0. To all Schroeder stimuli (F0=100Hz, SPL=30 dB, harmonics #1 - #400), there is a single continuous EPSP lacking “humps”. Like in Figure 7, A total of 100 representative traces are shown in each panel. **(B)** Voltage integrals from the same cell evoked by pure tones at 70 dB SPL. The triggering frequencies of the EPSPs in (A) are indicated in the graph and are part of the single-peaked frequency function. **(C)** Median voltages responses to different sound levels (20 – 50 dB SPL) from the same cell. Unlike the cell in Figure 6D, there are no clear “hump EPSPs”, suggesting that the inputs are distributed smoothly across frequencies. The EPSPs widen with increasing sound level, likely reflecting recruitment of additional inputs. **(D)** Halfwidths of the EPSPs in (C) shows widening with sound level at all curvatures, including C=0.

